# Inclusion of the ζ-chain drives phosphotyrosine signalling in CD19-CAR T cells

**DOI:** 10.1101/2025.07.04.663030

**Authors:** Aurora Callahan, Ryan Z Puterbaugh, Timothy Ro, Xinyan Zhang, Xiaolei Su, Arthur R Salomon

## Abstract

Although chimeric antigen receptor (CAR) T cell therapy has revolutionised individualised cancer therapies for relapsed/refractory lymphomas, signalling mechanisms underlying CAR T activation remain incompletely understood, especially among the three generations of CAR T exploiting different signalling domains. Here, using Jurkat T cell as a model, we investigate how costimulation influences tyrosine phosphorylation cascades using LC-MS/MS based phosphotyrosine (pY) proteomics and CD69 expression in the presence of small molecule inhibitors of key TCR signalling regulators. We find that including the ζ-chain in first (ζ-CAR), second (28ζ-CAR and BBζ-CAR), and third (28BBζ-CAR) generation CARs largely determines pY signalling, irrespective of costimulation. Further, we show that the phosphatase activity of PTPN22 and SHP-1 are largely negligible for activation of CARs, but indiscriminate inhibition of phosphatases using Pervanadate (PV) selectively activates BBζ-CARs without antigen encounter. Finally, we find that selective, partial inhibition of Itk using Soquelitinib reduces basal CD69 expression in Jurkat CAR T cells while maintaining their ability to activate in response to antigen. Our data suggest that the ζ-chain determines the pY signalling profile of CD19-CAR Jurkat T cells and that Itk may drive antigen-independent CD19-CAR activation.

## 1 Introduction

Chimeric antigen receptor (CAR) T cell therapy is a monumental achievement in the advancement of immunotherapies, providing the first potential modular therapeutic for a variety of disease states including autoimmune disorders and cancers(*1–3*). CAR designs include an extracellular targeting domain, typically a single chain variable fragment (scFv) of a highly specific antibody, a transmembrane domain, typically from T cell protein CD8 or CD28, and one or more intracellular signalling domains scavenged from the T cell receptor (TCR) itself and/or costimulatory receptors(*2*, *4*). In combination, the structural components of CARs are designed to ligate a disease associated antigen (DAA), promote immune synapse formation(*5–7*), initiate T cell activation through canonical TCR signalling networks(*8–10*), and mount an immune response towards the targeted cell(*11*). Notably, high antigen density on tumours and low antigen density on healthy cells is a requirement for the safety and efficacy of CAR T cells(*12*, *13*), leading some researchers to propose split-CARs that only promote T cell activation in the presence of two antigens expressed on the target cell(*14*, *15*). Regardless of design, the promise of a treatment adaptable to a specific disease state is highly desirable and has the potential to reshape personalised medicine.

Chimeric antigen receptors are designed to engage a TCR-like signalling cascade that results in cytokine expression and granzyme/perforin release. Numerous groups have evaluated the signalling capacity of CARs using various low throughput(*16*, *17*) and high throughput(*8–10*, *18*) methods and proposed similar models for CAR-based T cell activation. Work from Salter *et al.* (2018), Salter *et al*. (2021), and Griffith *et al*. (2022) used mass spectrometry based phosphotyrosine (pY) proteomics to show that primary CAR T cells or model Jurkat CAR T cells (CAR-Jurkats) induce phosphorylation changes similar to those of canonical TCR signalling with some key differences(*8*, *10*). First, Lck associated with CD28 costimulated CARs phosphorylates TCRζ ITAMs of high molecular weight (i.e., on the CAR), promoting the binding of Zap70 to the CARs TCRζ ITAMs and subsequent activation of Zap70 by Lck.

Activated Zap70 and Lck engage the LAT-SLP76 signalosome, although LAT itself is dispensable for CAR activation in third generation CD28 and 4-1BB costimulated CARs(*5*). Studies frequently observe phosphorylation of Itk^Y512^ and Itk substrates PLCγ1^Y783^ and GADS^Y45^, which are thought to act as signal integrators for CD28 costimulation(*19*). Notably, CD3ε/CD3γ/CD3δ ITAMs are seldom observed as significantly induced after CAR engagement, and further low throughout investigation has shown that CAR- and TCR-based signalling cascades are independent(*9*, *20*). Despite the potential similarities in pY signalling, costimulation of CARs can have profound effects on constitutive activation, retention, and exhaustion(*21*, *22*), suggesting that there exist unexplored activation pathways in CAR T cells with different costimulatory domains.

Recent approaches to improve CAR T cell therapy include screening hundreds of CAR designs in model systems for repeatable T cell activation, high CAR retention, and reduced tonic T cell activation. Many researchers use the Jurkat model T cell line during CAR screening(*23–26*) or for basic research on CAR-T cell activation due to its ease of modification, heavily studied T cell activation mechanisms, and speed with which CAR function can be assessed(*27*). Recently, Jahan *et al*. (2023) used a CAR-Jurkat model to evaluate the role of spacer length preceding the scFv region of CARs and found that CAR-Jurkats recapitulate many phenotypes of primary cells, including cytotoxicity, NF-κB/NFAT engagement, and intracellular phosphosignalling(*28*).

Similarly, Wan *et al.* (2025) used a CAR-Jurkat model to show that SMAD4 is involved in regulating Lck expression and, therefore, tonic expression of CD69 and CAR T cell proliferation(*29*). Finally, CAR-Jurkats were recently used by Gordon *et al.* (2022) to develop the CARPOOL library, which aims to determine an optimal set of intracellular signalling domains by barcoding hundreds of ICD combinations, expressing them in Jurkats, and assessing their activation potential(*26*). Despite the many drawbacks of clonal cell lines for biomedical research(*30*), the use of CAR-Jurkats has provided a platform for evaluating new CARs and the mechanism of CAR T cell activation that is demonstrably useful for CAR engineering.

To clarify the use of Jurkat T cells for CAR screening and research and determine the effects of key TCR feedback mechanisms in the context of CAR T cell activation, we leveraged our previously described SILAC co-culture pY mass spectrometry methodology(*10*) in tandem with CD69 flow cytometry assays. Using variable minimised first, second, and third generation CAR constructs mimicking the therapeutic Yescarta(*31*), we find that CAR-induced pY signalling is dominated by inclusion of TCRζ. Further, we show that all CARs show higher CD69 positive populations compared to non-transfected Jurkat T cells, but costimulation is required for a large-scale induction of CD69 expression. We also observe that CD19-CAR Jurkats differ from primary CAR T cell studies in that SHP-1 phosphatase activity shows no influence on CAR activation, but phosphatase activity globally is crucial for thresholding T cell activation. Finally, we find that pharmacological inhibition of Itk reduces tonic CD69 expression while maintaining co-culture induced activation. Overall, we provide a thorough evaluation of CAR signalling in Jurkat T cells and highlight the importance of Itk for basal activation in ζ-containing CD19-CAR Jurkats.

## 2 Materials and Methods

### 2.1 Cell culture

Jurkat T cells and CD19-CAR expressing Jurkat T cells, Raji B cells, J.muOT1.hCD8 (J^OT1^), and T2-K^b^ cells were maintained in RPMI 1640 supplemented with 10% FBS (Peak Serum #PS-FB3 Lot 01Z2233), 100 u/mL penicillin, 2 mM L-glutamine, 100 ug/mL streptomycin (100× PSQ, Cytiva #SV30082.01), and 2.5 ug/mL Plasmocin (InvivoGen #ant-mpp) in a humidified incubator at 37*C and 5% CO_2_. Chimeric antigen receptor-expressing Jurkat T cells were transduced as previously described(*32*). For stable isotopic labelling of amino acids in cell culture (SILAC) co-culture experiments, Jurkat T cells were maintained in SILAC RPMI 1640 (Thermo #A33823) supplemented with 10% dialysed FBS, 100 u/mL penicillin, 2 mM L-glutamine, 100 ug/mL streptomycin, 2.5 ug/mL Plasmocin, 0.28 mM ^13^C_6 15_N_4_ Arginine (Sigma Aldrich #608003-4.00G) and 0.22 mM ^13^C_6 15_N_2_ Lysine (Cambridge Isotope Laboratories #CNLM-291-H-1) for seven days.

### 2.2 Inhibitor treatment and stimulation

For inhibitor treatment experiments, Jurkat T cells were collected, washed once in plain RPMI 1640, then resuspended in the inhibitor solutions - 0.2% DMSO, 5 μM PP1 (MedChemExpress #HY-13804-5mg), 2 μM Soquelitinib (MedChemExpress #HY-150298), 5 μM U0126 (Cell Signaling Technologies #9903), 4 μM TPI-1 (MedChemExpress #HY-100463-5mg), 2 μM PTPN22-IN-1 (MedChemExpress #HY-139693-5mg), 10 μM Pervanadate - at a concentration of 3×10^7^ cells/mL for 3 hours at 37 C, 5% CO_2_ in a humidified incubator. After pretreatment, cells were washed once in 1× DPBS (with inhibitor), resuspended in inhibitor DPBS at a concentration of 2×10^8^ cells per millilitre, before inducing stimulation by crosslinking 2 μg/mL α-CD3ε (clone OKT3, Invitrogen #16-0037-85) and 2 μg/mL α-CD28 (clone CD28.6, Invitrogen #16-0288-85) with 22 μg/mL α-Mouse IgG (Jackson ImmunoResearch #115-005-062) secondary antibody. Cells were lysed after 5 minutes of stimulation by 1-to-1 addition of Urea Lysis Buffer (8 M Urea, 1 μM sodium orthovanadate, 1 μM β-glycerophosphate, 1 μM sodium pyrophosphate) before briefly sonicating at 70% amplitude for 15 second and dilution in 2× Lamelli Buffer (4% SDS, 125 mM TRIS-HCl pH 6.8, 20% glycerol, 5% β-mercaptoethanol, 0.01% bromophenol blue). Samples were stored at -20 C until use.

### 2.3 Flow cytometry

To evaluate CAR expression, one million CAR-expressing Jurkat T cells were collected and washed twice in FACS buffer (2% FBS in 1× DPBS), then fixed with 2% paraformaldehyde (PFA) at RT in the dark. After fixation, the cells were washed thrice with FACS buffer and stored before analysis on a Cytek Aurora flow cytometer. Analysis of expression was performed using Floreada and raw counts were visualised using Python 3.

For evaluating CD69 expression before and after co-culture, Jurkat T cells and CD19-CARs were pretreated with inhibitor for 1 hour as described in Section 2.2. For co-cultures, Raji B cells were fixed by 1:1 addition of PFA at room temperature for 15 minutes. After pretreatment with inhibitor, Jurkat cells were washed once in complete RPMI 1640 with inhibitor, then incubated at a concentration of 5×10^6^ T cells/mL in complete RPMI 1640 with inhibitor. For tonic signalling assays, Jurkats were maintained alone and for co-culture activation assays, Jurkats were maintained with fixed Raji B cells (pipetting up and down to mix) for 24 hours at 37 C, 5% CO_2_ in a humidified incubator. After 24 hours of incubation, cells were collected and resuspended in FACS buffer before staining with α-CD3ε PE (BD Biosciences #555333, for Jurkats) and/or α-CD69 APC (BD Biosciences #560967, for all samples). Samples were washed 3 times with 1 mL of FACS buffer, then fixed by 1-to-1 addition of 4% PFA for 15 minutes at room temperature, washed 3 times in 1 mL FACS buffer, then stored in FACS buffer until analysis on a Cytek Aurora flow cytometer. Analysis of percent CD69 positive cells was performed using Floreada and statistical analysis was performed using Python3.

### 2.4 Co-culture stimulation

Co-culture sample preparation for Western blotting or mass spectrometry sample preparation was performed as previously described(*32*). Briefly, Jurkat T cells, J^OT1^ cells or CD19-CARs and live Raji B cells were washed once in 1× DPBS, then resuspended at a concentration of 2×10^8^ (T cells) or 1×10^8^ (target cells). T2-K^b^ cells were loaded with OVA^257-264^ peptide by washing once in plain RPMI 1640, then incubating in plain RPMI 1640 supplemented with 1 μM OVA^257-264^ for 3 hours at 37 C, 5% CO2 in a humidified incubator. After incubation, the peptide-loaded T2-K^b^ cells (pT2-K^b^) were collected and washed once in 1× DPBS, then resuspended at a concentration of 1×10^8^ cells/mL. Cells were co-cultured by 1:1 (v/v, 2:1 E:T) addition of T cells to target cells, briefly centrifuged for 30 seconds at 500 ×g, incubated at 37 C for the indicated times, then lysed with Urea Lysis Buffer and cooled on ice for at least 15 minutes. Samples were then sonicated at 70% amplitude for 15 seconds (Western blotting samples) or 30 seconds (mass spectrometry samples). Samples for Western blotting were diluted 1:1 in 2× Lamelli buffer (4% SDS, 125 mM TRIS-HCl pH 6.8, 20% glycerol, 5% β-mercaptoethanol, 0.01% bromophenol blue), boiled at 95 C for 5 minutes, then stored at -20 C until use.

### 2.5 Western blotting

Western blotting was performed as previously described(*32*). Ten percent Tris-Glycine gels were poured in house and samples were separated at 100 V for 130 minutes before transferring to a PVDF membrane (EMD Millipore #IPFL00010) at 100 V for 100 minutes in an ice cold wet transfer apparatus. Membranes were blocked with Intercept TBS blocking buffer (LI-COR #927-60001) that was diluted 1 to 1 in 1× TBS, then incubated with primary antibody solution (antibody in 5% BSA dissolved in 1× TBST) overnight with gentle shaking. Membranes were washed 3× in 1× TBST before incubating in secondary antibody solution for 1 hour at room temperature on a rocker. Finally, membranes were washed 3× in 1× TBST, 2× in 1× TBS, then imaged on a LI-COR Odyssey CLx or LI-COR Odyssey M scanner. Western blot quantification was performed as previously described(*32*, *33*).

Primary Antibodies: PLCγ1 (1:10000, Millipore Sigma #05-163), PLCγ1^Y783^ (1:2000, Cell Signalling Technologies #2821), Erk1/2 (Cell Signalling Technologies #9107), Erk^T202Y204^ (1:2000, Cell Signalling Technologies #9101), Lck (1:4000, Cell Signaling Technologies #2657), Src family kinase activation site (Src^Y416^; 1:2000, Cell Signaling Technologies #2101).

Secondary Antibodies: IRDye 680RD Donkey anti Rabbit IgG (LI-COR #926-68073), IRDye 800CW Donkey anti Rabbit IgG (LI-COR #926-32213)

### 2.6 Sample preparation for LC-MS/MS

Mass spectrometry samples were prepared as previously described(*32*). Briefly, Urea-lysed samples were protein-input normalised (15 mg per sample for Jurkat, ζ-CAR, 28ζ-CAR, BBζ-CAR, 28BBζ-CAR and 10 mg per sample for J^OT1^ pT2-K^b^) by BCA (ThermoThermo #23225), adjusted to 2 mL with 4 M Urea Lysis Buffer, then diluted with 2.45 mL of 20 mM HEPES. Proteins were reduced with 50 μL of 1 M dithiothreitol (final concentration 10 mM) then alkylated with 500 μL of 100 mM iodoacetamide (final concentration 10 mM) before overnight digestion with trypsin (Promega #V5113) at a 1:100 trypsin:protein ratio. The following day, samples were acidified to 1% trifluoroacetic acid and desalted using Waters Sep-Pak C18 vacuum cartridges (Waters #WAT020515) as previously described(*10*). For pY peptide enrichment, the Src SH2 superbinder (sSH2) method was used as previously described(*10*, *32*, *34*).

### 2.7 LC-MS/MS, database searching, and analysis

Enriched pY peptides were separated on a Vanquish Neo uHPLC using a trap and elute setup. Peptides were first loaded onto a 7 cm Acclaim PepMap C18 trap column (2 cm bed length, 3 mm particle size, 100 Å; Thermo #164946) before elution and separation using an Aurora Ultimate C18 column (25 cm × 75 mm ID, 1.7 mm particle size; IonOpticks #AUR3-25075C18) using a 60 minute gradient at a constant 300 nL/minute starting at 5% Solvent B (80% acetonitrile, 0.1% formic acid) and 95% Solvent A (0.1% formic acid) and ending at 30% Solvent B and 70% Solvent A over a course of 50 minutes, then spiking to 95% Solvent B for 5 minutes and 99% Solvent B for the remaining 5 minutes. After each sample, the trap column was washed using a dedicated 60 minute method consisting of 12 zebra washes and the analytical column was washed with 3 aqueous/organic cycles before equilibrating for the next sample. In between each set of 5 samples, 20 μL of Flush solution (Thermo #) was injected, and 1 μg of BSA was run to assess instrument sensitivity.

Data was collected on an orbitrap Ascend mass spectrometer. For MS settings, the spray voltage was set to 2050 V, the ion transfer tube was set to 275 C, and a lock mass of 445.12003 was used. The orbitrap detector was used for MS1 scans with wide quad isolation, a resolution of 120,000, a mass range of 300-1500 m/z, a maximum injection time of 50 ms, 250% normalised AGC target, and RF lens at 30% in centroid mode. The following filters were included: monoisotopic peak and peptide mode, charge state 2-7, 375-1500 m/z precursor selection range, dynamic exclusion (exclude after 1 times, 15 second exclusion time, 10 ppm low and high mass tolerances, excluding isotopes), intensity range of 20,000 (min) - 1E20 (max) with relative intensity threshold of 20%. MS2 were collected in data-dependent acquisition mode using a 3 second cycle time, 30% normalised higher-energy collisional dissociation (HCD), and a quadrupole isolation width of 0.7 m/z. Fragment ions were collected in centroid mode and detected using the ion trap in Turbo Mode with a scan range of 150-2000 m/z with a maximum injection time of 35 ms, 100% normalised AGC, and using the dynamic mode for maximum injection time.

Thermo .RAW files containing MS spectral data were analysed using the high throughput autonomous proteomics pipeline (HTAPP) and PeptideDepot as previously described(*35*, *36*). Briefly, MS2 spectra were searched against the UniProt complete proteome dataset (downloaded August 2019 with 98,300 unique forward sequences) using the Mascot search engine to 0.1% FDR using a reverse decoy database strategy. For peptide spectrum matching, the following parameters were used; trypsin enzyme cleavage with up to 2 missed cleavages, 7 ppm precursor mass tolerance, 500 mmu fragment ion mass tolerance, variable modifications for phosphorylation (S/T/Y +79.9963 Da) and oxidation (M +15.9949 Da), with static modifications of carbamamidomethylation (C +57.0215 Da). Retention time alignment and integration of selected ion chromatograms was used to determine the relative peptide abundance for each PSM as previously described(*37*). The resulting peak areas were further processed in PeptideDepot to normalise peak areas to a spike in pY peptide (LIEDpYTAK; samples #44 and #75 were dropped due to no sequencing of this peptide), then statistical significance between log2 mean abundance for the time points was determined using Welch’s T test and FDR was estimated using the method of Storey(*38*). Replicate reproducibility was determined with multiple linear regression and principal component analysis. Post-translational modification signature enrichment analysis was performed as previously described(*32*, *39*) using the R-script ssGSEA2.0.R with the PTM-signature database version 2.0.0.

### 2.8 Data availability

All plotting was performed using Python3.12.5 in a JupyterLab environment with the dependencies matplotlib (version 3.9.1), scipy (version 1.14.0), numpy (version 2.0.1), scikit-learn (version 1.5.1), pandas (version 2.2.2), and a custom ‘helpers’ module used for beautifying matplotlib graphs. Statistical analysis of Western blots and flow cytometry data was performed using a self-coded version of the Holm-Sidak FWER correction based on the algorithm provided by GraphPad Prism. All Image Studio files (Westerns), post-gating flow cytometry .csv files, and code used in this project are available in the supporting materials and on GitHub (https://github.com/Aurdeegz/CAR-Generations). Mass spectrometry .RAW files are available from the ProteomeXchange Consortium via the PRIDE repository (Dataset ID: PXD065669, Reviewer Username, Reviewer Password: DHakF6gr0r7v).

## 3 Results and Discussion

### 3.1 CD19-CAR expression and mass spectrometry experimental design

Because of the large clinical variation in CAR function between CARs with different costimulatory domains, we seek to evaluate the pY signalling capacity of CD19-CAR T cells with clinically relevant costimulatory domains modelled after the therapeutic Yescarta (Axicabtagene ciloleucel)(*31*). We engineer Jurkat T cells to express to express a first generation (ζ; ζ-CAR hereafter), two second generation (CD28co-ζ; 28ζ-CAR hereafter, 41BBco-ζ; BBζ-CAR hereafter), and one third generation (CD28co-41BBco-ζ; 28BBζ-CAR hereafter) targeting CD19 with an FMC63-based scFv, CD8 transmembrane domain and hinge region, and an intracellular superfold GFP tag (Figure 1A, Supporting Figure 1). All CD19-CARs are highly expressed compared with a Jurkat T cell control, with little variation in the GFP+ population between CD19-CAR constructs (Figure 1B). Co-culture between CD19-CARs and Raji B cells show significant induction of Erk1^T202Y204^/Erk2^T185Y187^ and PLCγ1^Y783^ at 5 and 15 minutes post co-culture, and 28BBζ-CAR shows significantly lower levels of basal Erk1^T202Y204^/Erk2^T185Y187^ and PLCγ1^Y783^ phosphorylation than WT Jurkats (Figures 1C-E, Supporting Figure 2, Supporting Table 1). It is important to note that Raji B cells also express Erk1/2 and PLCγ1, and thus the basal levels of Erk1^T202Y204^/Erk2^T185Y187^ and PLCγ1^Y783^ may be influenced by Raji phosphorylation levels. However, we have previously shown that Raji B cells display little-to-no pY changes after co-culture(*10*, *32*), thus the induction we observe is most likely due to the presence of CD19-CARs. To gain precise information about the pY signalling potential, we adopt a co-culture Western blot and phosphoproteomics workflow as described previously(*10*, *32*). Briefly, CD19-CAR T cells and Raji B cells are cultured to confluence in standard RPMI and ^13^C_6 15_N_4_ Arginine/^13^C_6 15_N_2_ Lysine RPMI, respectively, before mixing at a 2:1 effector-to-target (E:T) ratio, briefly centrifuged to promote contact, and allowed to incubate for 0 minutes, 5 minutes, or 15 minutes before lysis in 8 M Urea. A portion of the lysate is saved for phosphosite specific Western blotting, and the remaining lysate (15 mg protein per sample) is processed through a standard bottom-up proteomics workflow(*10*, *32*, *37*). After, pY peptides are enriched using the Src SH2 superbinder domain(*10*, *32*, *34*) and analysed by LC-MS/MS on an Orbitrap Ascend mass spectrometer. These methods provide precise information regarding the pY signalling cascades initiated by each CAR during co-culture.

**Figure 1:**
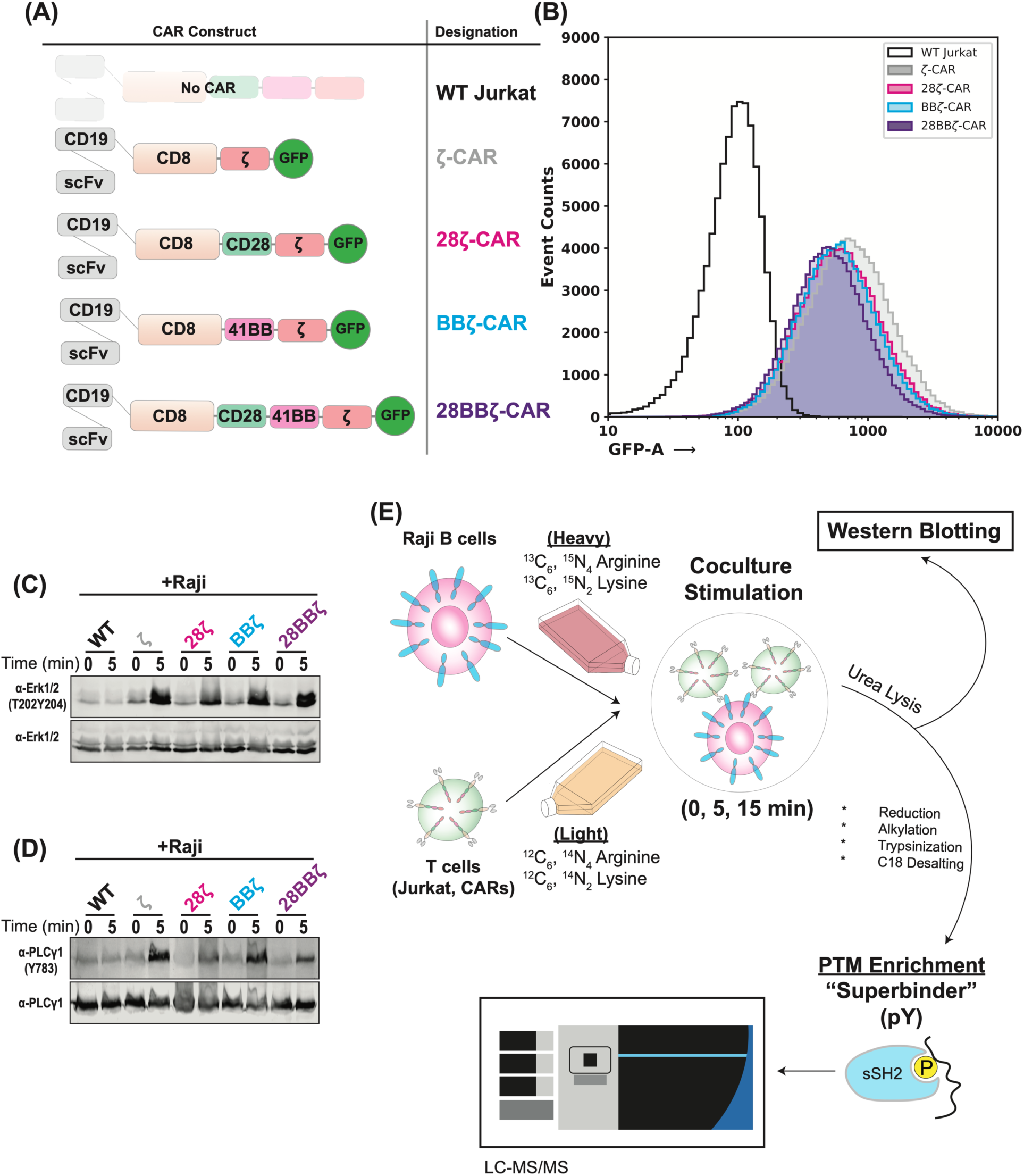
Expression of CD19-CARs and phosphoproteomic workflow. (A) Schematic representation of the cell lines used in this study. (B) Flow cytometry analysis showing total GFP expression in CD19-CAR Jurkats. (C-D) Western blot analysis of co-culture samples showing phosphorylation of Erk1/2 and PLCγ1, respectively. (E) Flow chart depicting the co-culture stimulation phosphoproteomics workflow used in this study.

**Figure 2:**
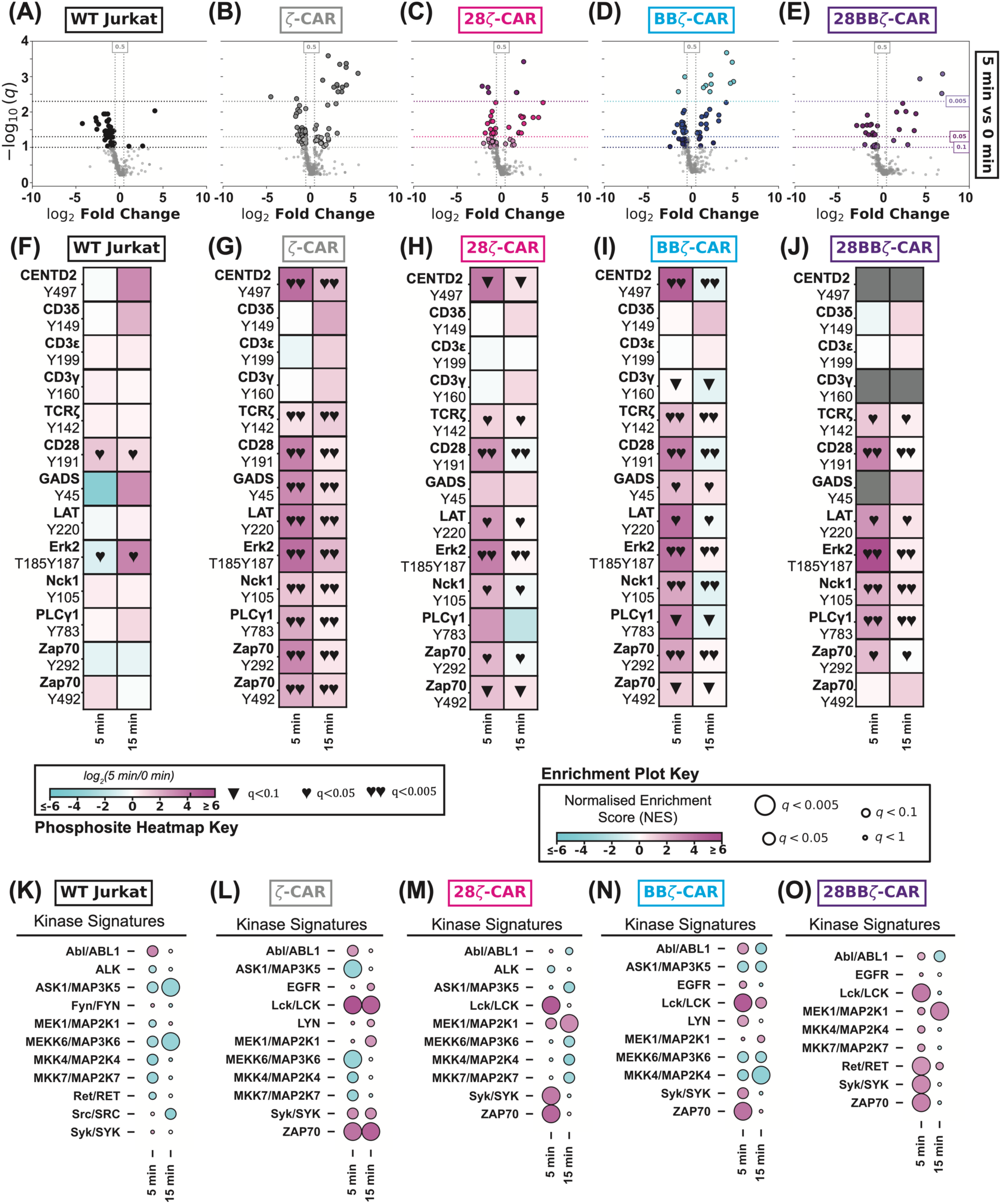
Inclusion of the ζ-chain in CD19-CARs dominates pY signalling. (A-E) Volcano plot comparing unique pY sites observed from 5 minutes and 0 minutes of co-culture between Raji cells and WT Jurkats, ζ-CAR, 28ζ-CAR, BBζ-CAR, and 28BBζ-CAR, respectively. (F-J) Heat maps comparing 5 minutes or 15 minutes with 0 minutes of co-culture between Raji cells and WT Jurkats, ζ-CAR, 28ζ-CAR, BBζ-CAR, and 28BBζ-CAR, respectively. Sites presented are a selection of known TCR-responsive pY sites. ▼ indicates q < 0.1 (trending towards significance), ♥ indicates q < 0.05, and ♥♥ indicates q < 0.005. (K-O) Post translational modification signature enrichment analysis using pY site data showing enrichment of PhosphoSitePlus kinase substrate signatures for WT Jurkats, ζ-CAR, 28ζ-CAR, BBζ-CAR, and 28BBζ-CAR, respectively.

### 3.2 Inclusion of the ζ-chain in CD19-CARs dominates pY signalling cascades

Using SILAC co-cultures with pY enrichment mass spectrometry (Figure 1C), we sequence a total of 496 pY sites on 340 unique proteins across all samples with high reproducibility (R>0.8) and replicate clustering by principal component analysis (Supporting Figure 4, Supporting Tables 2-7). In line with our own previous work(*10*, *32*), we observe few pY sites significantly elevating in co-cultures of Jurkat and Raji B cells, and many pY sites significantly increasing in abundance after co-culture of CD19-CARs with Raji B cells (Figure 2A-E). Notably, the pattern of significantly changing pY sites after 5 minutes of co-culture closely matches co-cultures of Jurkat cells expressing the murine OT1 transgenic TCR and CD8 (J^OT1^) co-cultured with T2-K^b^ cells expressing a class I MHC loaded with OVA^257-264^ (pT2-K^b^; Supporting Figures 3C-D, 5A-B), which is a well established model for studying T cell signalling and T cell activation(*40–42*). Across all CD19-CARs, many of the significantly induced pY sites are involved in the forward progression of T cell signalling, including the immunotyrosine based activation motifs (ITAMs) ζ^Y123^ and ζ^Y142^, CD28^Y191^, GADS^Y45^, LAT^Y220^, Erk1^Y204^ and Erk2^Y187^, Nck^Y105^, PLCγ1^Y771^, PLCγ1^Y775^, PLCγ1^Y783^, PKCδ^Y313^, SHC1^Y427^, WASL^Y256^, Zap70^Y292^, Zap70^Y492^, and Zap70^Y493^ (Figure 2F-J). In contrast, co-cultures of JOT1 and pT2-K^b^ cells show significant phosphorylation of ITAMs on other components of the TCR, such as CD3δ^Y149,^ CD3δ^Y160^, CD3ε^Y188^, and CD3ε^Y190^, and regulatory sites on Zap70 such as Zap70^Y315^ and Zap70^Y319^ (Supporting Figure 5C), suggesting that CD19-CAR T cells engage a subset of the TCR signalling pathway that is independent of the TCR itself. Using post-translational modification signature analysis (PTM-SEA), a computational tool that discerns patterns between measured PTM proteomics data and annotated databases, shows downregulation of both kinase substrate signatures and pathways in co-cultures of WT Jurkats with Raji B cells (Figures 2K, Supporting Figure 6A). In contrast, all CD19-CARs show upregulation of kinase substrate signatures particularly the Src family kinase Lck and the Syk family kinase Zap70, which are both necessary for the forward progression of T cell activation through CARs and the TCR (Figures 2L-O)(*43*). Similarly, all CD19-CARs show upregulation of signalling pathways, generally, with ζ-CAR showing the weakest and 28BBζ-CAR showing the strongest engagement with signalling cascades (Supporting Figure 6B-E). Together, our data suggest that the inclusion of ζ-chain is the primary determinant of pY signalling in CD19-CAR Jurkats and that ζ-containing CARs engage only a subset of canonical TCR signalling.

**Figure 3:**
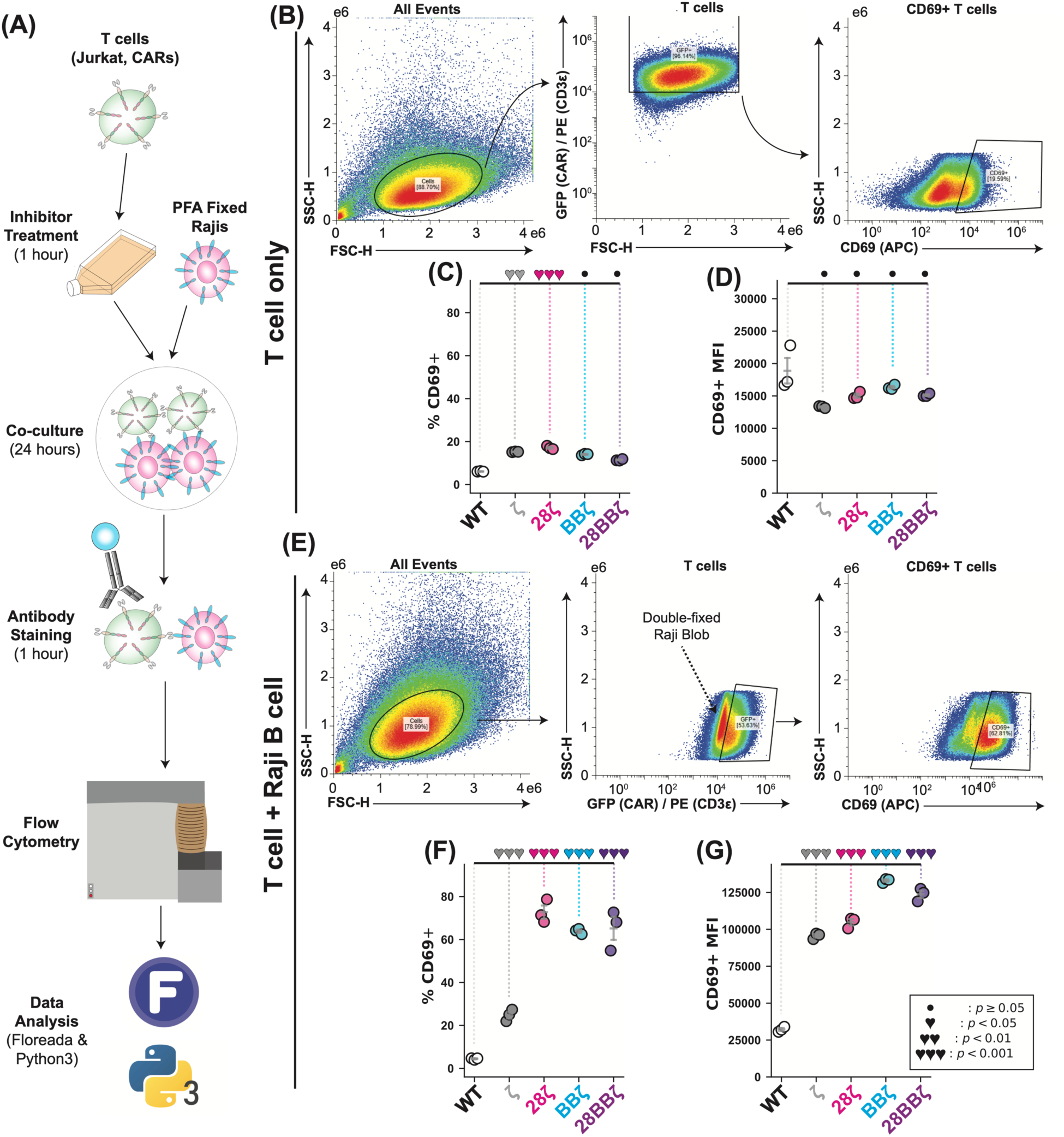
Costimulation is required for elevated CD69 expression after co-culture, however ζ-CAR Jurkats show tonic CD69 expression. (A) Flow chart showing the experimental design of CD69 activation assays. (B) Gating strategy for non co-culture CD69 flow cytometry data analysis. (C-D) Evaluation of percent positive CD69 cells and the mean fluorescence intensity of CD69 positive cells, respectively, for non co-cultured CAR T cells. (E) Gating strategy for co-culture CD69 flow cytometry data analysis. (F-G) As in C and D, except for co-culture samples.

**Figure 4:**
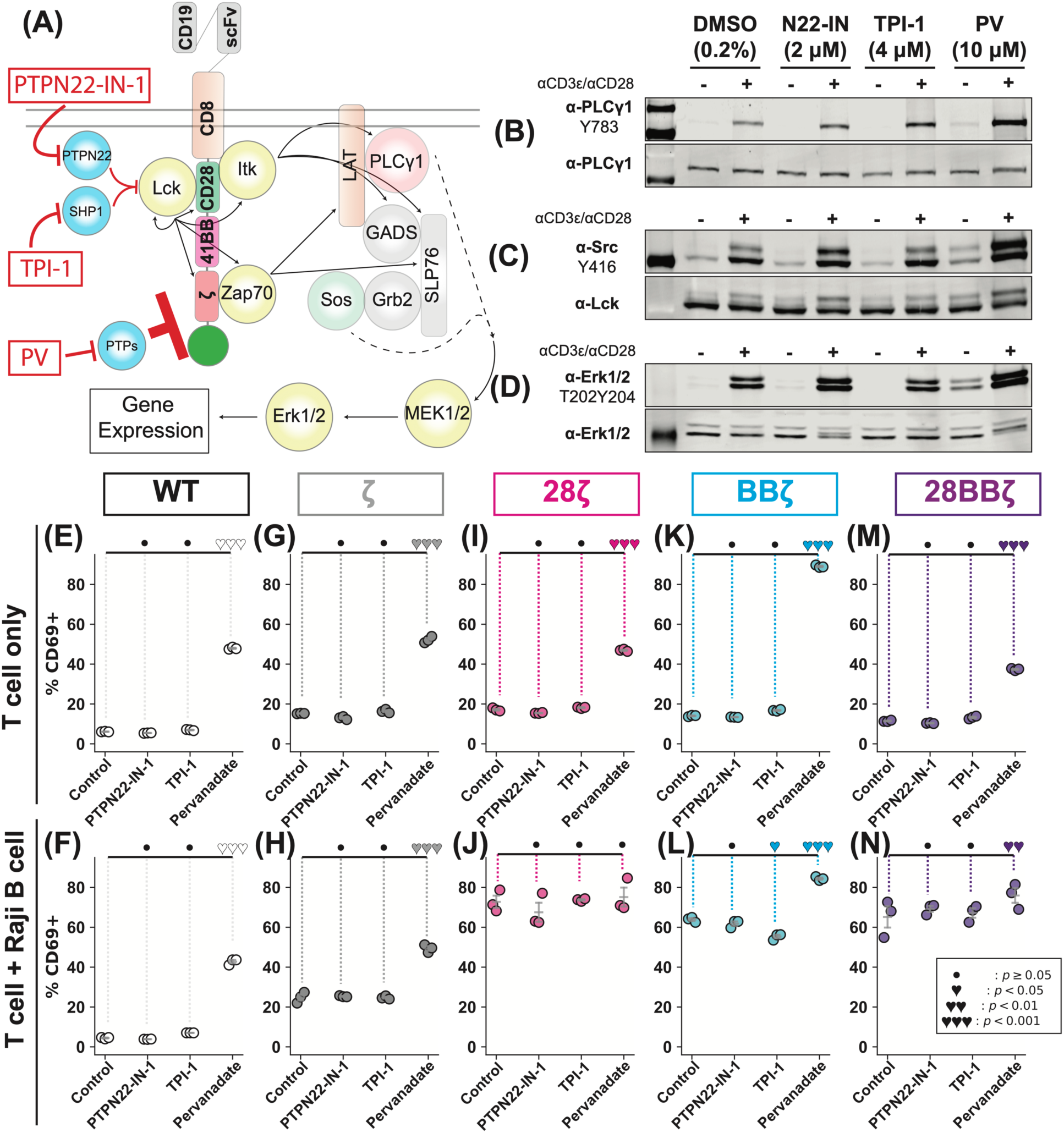
PTP activity globally, but not TCR-specific regulatory PTPs, restrict CD69 expression in a costimulation-dependent manner. (A) Simplified schematic representation of CAR T cell signalling with inhibitors and their targets noted. (B-D) Western blot analysis of Lck^Y394^, Zap70^Y493^, and Erk1^T202Y204^/Erk2^T185Y187^, respectively, in samples treated with PTPN22-IN-1, TPI-1, or PV. (E-F) Dotplot showing the effect of PTPN22-IN-1, TPI-1, and PV on the percentage of CD69 positive WT Jurkats in non co-culture and co-culture samples, respectively. (G-H) As in E and F, except for ζ-CAR Jurkats. (I-J) As in E and F, except for 28ζ-CAR Jurkats. (K-L) As in E and F, except for BBζ-CAR Jurkats. (M-N) As in E and F, except for 28BBζ-CAR Jurkats.

**Figure 5:**
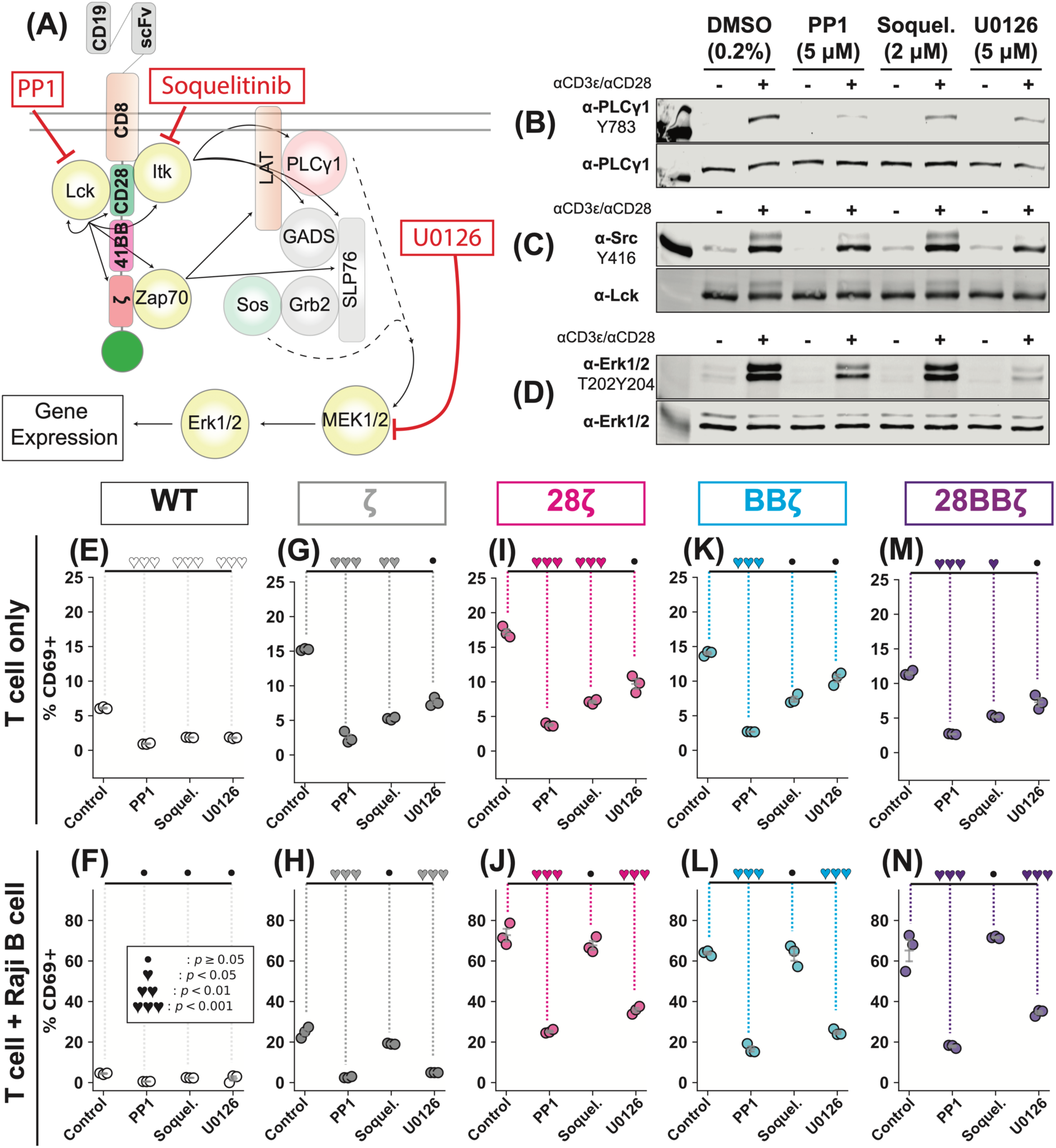
Pharmacological inhibition of Itk with Soquelitinib reduces tonic CD69 expression without impairing CAR T cell activation. (A) Simplified schematic representation of CAR T cell signalling with inhibitors and their targets noted. (B-D) Western blot analysis of Lck^Y394^, PLC1^Y783^, and Erk1^T202Y204^/Erk2^T185Y187^, respectively, in samples treated with PP1, Soquelitinib, or U0126. (E-F) Dotplot showing the effect of PTPN22-IN-1, TPI-1, and PV on the percentage of CD69 positive WT Jurkats in non co-culture and co-culture samples, respectively. (G-H) As in E and F, except for ζ-CAR Jurkats. (I-J) As in E and F, except for 28ζ-CAR Jurkats. (K-L) As in E and F, except for BBζ-CAR Jurkats. (M-N) As in E and F, except for 28BBζ-CAR Jurkats.

### 3.3 Inclusion of the ζ-chain in CD19-CARs increases basal CD69 expression but is not sufficient for activation after antigen encounter

Although CD19-CARs all effectively engage pY sites in the known T cell receptor signalling pathway, previous literature suggests that costimulation is necessary for T cell activation(*44*). To verify that costimulation is necessary for CAR T cell activation, we perform 24 hour co-culture stimulation experiments using PFA fixed Raji B cells and assess CD69 expression (Figure 3A), a known marker of T cell activation used in CAR-Jurkat screens(*25*, *26*), by flow cytometry. We assess tonic T cell activation by gating the main cell population, then gating on high signal in the GFP (CD19-CARs) or PE (α-CD3ε PE for WT Jurkats), then high signal for APC (α-CD69 APC) (Figure 3B). We observe a significant increase in percent CD69 positive cells in ζ-CAR and 28ζ-CAR, a trend toward significance in BBζ-CAR, and no significant increase in 28BBζ-CAR (Figure 3C), with no significant change in mean fluorescence intensity between CD69 positive WT Jurkats and any CD19-CAR (Figure 3D). To assess T cell activation during co-culture with PFA fixed Raji B cells, a similar gating strategy is used taking into account the large Raji B cell cluster that appears in GFP/PE around 10^4^ (Figure 3E). We again observe a significant increase in percent CD69 positive cells in ζ-CAR compared to WT Jurkat, however all costimulated CD19-CARs show significantly higher percent CD69 positive cell populations than WT Jurkat or ζ-CAR (Figure 3F, Supporting Table 8). In co-cultured samples, the mean fluorescence intensity of CD69 positive cells is significantly increased in all CD19-CARs (Figure 3G). Our data show that percent CD69 positive cells is a metric that reproduces known and consistently observed phenotypes for CD19-CAR T cells, particularly tonic CD69 expression.

### 3.4 Global PTP activity, but not PTPN22 or SHP-1, threshold CAR T cell activation

Phosphatases are quickly activated to coordinate the deactivation of T cells after T cell receptor ligation(*43*), however phenotypic differences in CAR T cell activation (e.g., exhaustion, tonic activation) suggest the potential dysregulation of these feedback mechanisms. To evaluate the role of PTPs in CAR T cell activation, we leverage small molecule inhibitors of the T cell PTPs PTPN22 (PTPN22-IN-1) and SHP-1 (TPI-1), as well as the pan-PTP inhibitor pervandate (PV). First, we validate the activity of 2 μM PTPN22-IN-1, 4 μM TPI-1, and 10 μM PV in Jurkat T cells using CD3ε and CD28 crosslinking to stimulate the TCR. While only PV treatment significantly elevates basal and stimulated Src family kinase activation site (SFK^Y416^), PLCγ1^Y783^, and Erk1^T202Y204^/Erk2^T185Y187^ abundance, TPI-1 and PTPN22-IN-1 treatment significantly elevate the SHP-1 substrate(***45***) PLCγ1^Y783^ and the PTPN22 pathway target(*46*, *47*) Erk1^T202Y204^/Erk2^T185Y187^, respectively (Figure 4B-D, Supporting Figure 7, Supporting Table 9), showing that these inhibitors function at the given concentrations. To investigate whether PTP activity influences CAR T cell activation, we treat CD19-CARs with each PTP inhibitor and assess CD69 expression after 24 hours of co-culture. In WT Jurkats and ζ-CAR, inhibition of PTPN22 or SHP-1 does not significantly alter percent CD69 positive cells in either basal or co-culture conditions, however treatment with 10 μM PV significantly increases percent CD69 positive cells in both basal and co-culture conditions (Figure 4E-H). Similarly, inhibition of PTPN22 or SHP-1 does not alter basal percent CD69 positive cells in 28ζ-CAR, BBζ-CAR, or 28BBζ-CAR and does not prevent co-culture induced increases in percent CD69 positive cells, while PV treatment uniformly elevates basal percent CD69 positive cells (Figure 4I-N). Notably, BBζ-CAR alone shows a significant reduction in percent CD69 positive cells when treated with the SHP-1 inhibitor TPI-1 during co-culture, and greater than 90% CD69 positive cells when treated with PV in the basal state (Figure 4K-L), suggesting CD19-CARs without CD28 costimulation are thresholded more by PTPs than CD28 costimulated CARs. Together, our results demonstrate that PTP activity globally is critical for regulating CD19-CAR activation, but the TCR-specific phosphatases PTPN22 and SHP-1 are not involved in thresholding CD19-CAR activation.

### 3.5 Inhibition of Itk reduces tonic CD69 expression while maintaining CD19-CAR T cell activation

The kinases Lck and Zap70, which are required for T cell activation through the TCR(*43*), play a major role in CAR T cell activation(*48*), however the role of other proximal and distal kinases, like Itk and MEK1/2, in CAR T cell activation remains unclear. To understand the role of kinases in CAR T cell activation, we first show that small molecule inhibitors of Lck (5 μM PP1), Itk (2 μM Soquelitinib), and MEK1/2 (5 μM U0126) function appropriately in Jurkat T cells. As expected, 5 μM PP1 and 5 μM U0126 significantly dampen TCR induced phosphorylation of PLC1^Y783^, Lck^Y394^, and Erk1^T202Y204^/Erk2^T185Y187^(*49*, *50*),while treatment with 2 μM Soquelitinib significantly reduces phosphorylation of the Itk substrate PLC1^Y783^ and pathway target Erk1^T202Y204^/Erk2^T185Y187^(*51*)(Figure 5B-D, Supporting Figure 8, Supporting Table 10). To determine whether these kinases perturb CAR T cell activation, we treat CD19-CARs with each kinase inhibitor and assess CD69 expression after 24 hours of co-culture. In WT Jurkats, we find that PP1, Soquelitinib, and U0126 reduce the percent CD69 positive cells in the basal state but not during co-culture (Figure 5E-F). In all CD19-CARs, PP1 treatment reduces tonic CD69 expression and PP1 or U0126 treatment significantly reduces CD69 expression after co-culture (Figures 5I-N). In agreement, Soquelitinib treatment significantly reduces percent CD69 positive cells in the basal state of 28ζ-CARs and 28BBζ-CARs while BBζ-CARs are trending toward significance (p = 0.06; Figures 5I, 5K, 5M). However, Soquelitinib-treated CD19-CARs are able to maintain the co-culture induced increase in percent CD69 positive cells at levels similar to untreated CD19-CARs (Figures 5J, 5L, 5N). Together, these data suggest that Itk acts to elevate basal levels of CD69 in CD19-CARs but is not required for activation induced CD69 expression.

## 4 Discussion

The development of novel chimeric antigen receptors with increased antigen sensitivity, specificity, and target clearance is necessary to improve the range of CAR T cell applications. Current methods for the identification of functional chimeric antigen receptors require high-throughput screening of CAR constructs expressed in model systems(*23–26*, *28*), such as Jurkat T cells, or low-throughput, logic driven construction(*9*, *52*). Using these systems, researchers have identified and verified novel CARs that are highly efficacious, outperforming the current clinically approved constructs in mouse models(*26*). However, Jurkat T cells are PTEN deficient and resulting in mislocalization of Itk(*53*), which each determine the fate of key TCR activation pathways. While the utility of the Jurkat model is empirically verified in the CAR field, a thorough evaluation of CAR Jurkat signalling in the context of costimulation, Jurkat TCR signalling, and key TCR protein activity has not been reported.

Here, we evaluate the signalling capacity of the CAR Jurkat model system using pY-based LC-MS/MS, activation assays, and small molecule inhibition of key TCR regulators. Our results demonstrate CAR signalling is highly similar, independent of costimulation, and very similar to pMHC/TCR ligation (Figures 1C-E, 2, Supporting Figures 2-5). We observe increases in TCRζ-ITAMs in CAR co-cultures but only TCR ligation induces phosphorylation of ITAMs on CD3ε, CD3γ, and CD3δ (Figure 2, Supporting Figure 5), supporting the literature precedent that CAR- and TCR-induced signalling have non-overlapping origins(*20*). Our results are in line with our own previous work, where we have demonstrated that CD19-28ζ-CAR Jurkats(*32*) and CD19-28BBζ-CAR Jurkats(*10*, *32*) very strongly engage a TCR-like signalling cascade. While there does not exist a variable minimised comparison with primary CAR T cells, studies evaluating CAR signalling in primary cells have demonstrated similar but notably different results. Salter *et al*. (2018) directly compared phosphorylation after activation of CD19-BBζ-CARs and CD19-28ζ-CARs in primary human T cells, finding that 28ζ-CARs more strongly engage the phosphoproteome than BBζ-CARs, however both engage pY sites in the TCR signalling pathway including phosphorylation of CD28^Y209^(*8*). In our work, we observe little difference in the strength of phosphoproteome engagement between 28ζ-CAR Jurkats and BBζ-CAR Jurkats, however BBζ-CAR Jurkats do engage CD28^Y191^ similarly to ζ-CAR, 28ζ-CAR, and 28BBζ-CAR (Figure 2). In a follow up study, Salter *et al*. (2021) directly compare TCR stimulation with MHC tetramer and 28ζ-CAR stimulation with activating antibodies in primary T cells, showing that TCR signalling is stronger than CAR signalling in a simplified stimulation model(*9*). Our results are in agreement, as OVA^257-264^ loaded T2-K^b^ cells strongly induce tyrosine phosphorylation along the TCR signalling pathway (Supporting Figure 5). Although our data and those of Salter *et al*. (2018) and Salter *et al*. (2021) have vastly different signalling kinetics and fold-changes, the different methods of stimulation (antigen/antibody coated beads versus centrifugal cell-to-cell contact) likely account for these differences. Overall, our results agree with the literature in that TCRζ containing CARs engage a similar TCR-like pathway, however the similar induction levels we observe may be an artefact of the Jurkat model system which need to be accounted for in screens for CAR function.

It is well established that costimulation can greatly affect the strength and timing of CAR T cell activation and whether CAR T cells exhaust or persist(*2*), which given our data and the literature is likely independent of early, forward progressing phosphorylation-based signalling. In our CAR Jurkat model, we also seek to determine whether early/late forward pathways or feedback are important for CAR-based T cell activation. Using a CD69 flow cytometry activation assay we show that CAR Jurkats require costimulation to increase the CD69 positive population, and that ζ-CAR and 28ζ-CAR show high basal CD69 populations (Figure 3). Using the CD69 activation assay with small molecule inhibitors of SHP-1 (TPI-1), PTPN22 (PTPN22-IN-1), and PTPs globally (PV), we find that the phosphatase activity of SHP-1 and PTPN22 is not important for regulating the activation of CD19-CARs, however global PTP activity thresholds CD69 expression particularly in BBζ-CARs (Figure 4). Previous literature has shown that PTPN22 deletion does not affect CAR clearance of solid tumours(*54*), however some literature suggests SHP-1 knockout enhances the killing ability of some CAR T cells(*55*) and SHP-1 negatively regulates CD19-BBζ-CAR activation(*56*). Further, T cell protein tyrosine phosphatase (TCPTP; gene name *PTPN2*) deletion has also been implicated in thresholding HER2-28ζ-CAR cytotoxicity(*57*). Our results demonstrating that SHP-1 phosphatase activity do not influence CD19-CAR activation, while global phosphatase inhibition does, suggests that negative regulation of CD19-CARs in Jurkats differs from CAR primary T cells and should be taken into account in CAR Jurkat screens.

Finally, we also investigate the role of feed-forward activation from CD19-CARs using small molecule inhibitors of Lck (PP1), Itk (Soquelitinib), and MEK1/2 (U0126). As expected, PP1 treatment ablates CD19-CAR-induced tonic CD69 expression and stunts antigen-induced activation, and inhibition of MEK1/2 acts similarly although to a lesser extent (Figure 5E-N). Interestingly, inhibition of Itk reduces the CD69 positive population to the level of untreated WT Jurkats in all CD19-CARs, with significant reductions in CD69 positive CAR T cells in ζ-CAR and 28ζ-CAR compared to untreated ζ-CAR and 28ζ-CAR (trending towards significance in BBζ-CAR; Figure 5E-N). Soquelitinib (also known as CPI-818) is a recently described selective, irreversible Itk inhibitor that was shown to reduce exhaustion markers and restore effector function to previously exhausted T cells(*51*), and was recently given an FDA Orphan Drug Designation for the treatment of T cell lymphomas. Soqeulitinib was recently used to show that the kinase activity determines the differentiation of naive CD4+ T cells into T helper 17 cells in a calcium dependent, MAPK independent manner(*58*), further demonstrating the unique role of Itk in T cell maintenance. Our results, showing that Itk inhibition can reduce tonic CD69 expression without impairing antigen-induced T cell activation irrespective of costimulation, provides a potential mechanism for reducing antigen-independent effects of CAR T cell therapies.

## 5 Conclusion

The rush to develop new chimeric antigen receptors with high specificity and low off-target effects has led to the use of classical T cell model systems for their reliability and well characterised activation pathways. Our results suggest that activation in CAR Jurkat T cells is dominated by TCRζ on the CARs at the pY level, but are only able to fully activate with costimulation. Further, we find that PTPs generally threshold CAR T cell activation, but PTPN22 and SHP-1 phosphatase activity in particular are not important for CAR Jurkat regulation.

Finally, we provide evidence that Itk inhibition can reduce tonic activation of CAR Jurkats without compromising antigen-induced activation, unlike inhibition of Lck or MEK1/2. Our findings show some limitations of the CAR Jurkat model and provide an avenue for combination therapies, namely inhibition of Itk, that could reduce adverse effects of tonic CAR T cell activation.

## Author Contributions

Conceptualisation: AC

Methodology: AC

Investigation: AC, RZP, TR, XZ

Visualisation: AC

Funding acquisition: ARS, XS

Project administration: AC, ARS, XS

Supervision: AC

Writing – original draft: AC

Writing – review & editing: AC, ARS, XS

## ACKNOWLEDGMENT

We would like to thank Dr. Nicholas DaSilva at the Brown University Proteomics Core facility for helping us operate the mass spectrometer, Dr. Kevin Carlson at the Brown University Flow Cytometry and Cell Sorting Facility for running the flow samples, and Dr. Christoph Schorl at the Brown University Genomics Core Facility for allowing us to use the LI-COR scanner for Western blotting. Funding Sources: NIH grant P01AI091580 (ARS), 1S10OD036295 (ARS), R35 GM138299(XS), R21 CA286364(XS), R21 CA294038(XS), NIH Director’s Transformative Research Award EB037112(XS), American Cancer Society Research Scholar Grant 135926(XS), Gabrielle’s Angel Foundation Medical Research Award(XS), Pershing Square Sohn Prize for Young Investigators in Cancer research(XS), Human Frontier Science Program Early-Career Research Grant RGY0088/2021(XS).

## ABBREVIATIONS

CAR: Chimeric antigen receptor
TCR: T cell receptor
scFv: single-fold variable fragment
DAA: disease associated antigen
pY: phosphotyrosine
Lck: Lymphocyte cytosolic kinase
Itk: Interleukin 2 inducible T cell kinase

## References

1. S. Abbasi, M. A. Totmaj, M. Abbasi, S. Hajazimian, P. Goleij, J. Behroozi, B. Shademan, A. Isazadeh, B. Baradaran, Chimeric antigen receptor T (CAR-T) cells: Novel cell therapy for hematological malignancies. Cancer Med. 12, 7844–7858 (2023).

2. S. E. Lindner, S. M. Johnson, C. E. Brown, L. D. Wang, Chimeric antigen receptor signaling: Functional consequences and design implications. Sci. Adv. 6, eaaz3223 (2020).

3. T. A. da Silva, CAR T Cell-Therapy for Infectious Diseases with Emphasis on Invasive Fungal Infections. Ther. Deliv. 12, 627–630 (2021).

4. M. Sadelain, R. Brentjens, I. Rivière, The Basic Principles of Chimeric Antigen Receptor Design. Cancer Discov. 3, 388–398 (2013).

5. R. Dong, K. A. Libby, F. Blaeschke, W. Fuchs, A. Marson, R. D. Vale, X. Su, Rewired signaling network in T cells expressing the chimeric antigen receptor (CAR). EMBO J. 39, e104730 (2020).

6. A. J. Davenport, R. S. Cross, K. A. Watson, Y. Liao, W. Shi, H. M. Prince, P. A. Beavis, J. A. Trapani, M. H. Kershaw, D. S. Ritchie, P. K. Darcy, P. J. Neeson, M. R. Jenkins, Chimeric antigen receptor T cells form nonclassical and potent immune synapses driving rapid cytotoxicity. Proc. Natl. Acad. Sci. 115, E2068–E2076 (2018).

7. P. J. Chockley, J. Ibanez-Vega, G. Krenciute, L. J. Talbot, S. Gottschalk, Synapse-tuned CARs enhance immune cell anti-tumor activity. Nat. Biotechnol. 41, 1434–1445 (2023).

8. A. I. Salter, R. G. Ivey, J. J. Kennedy, V. Voillet, A. Rajan, E. J. Alderman, U. J. Voytovich, C. Lin, D. Sommermeyer, L. Liu, J. R. Whiteaker, R. Gottardo, A. G. Paulovich, S. R. Riddell, Phosphoproteomic analysis of chimeric antigen receptor signaling reveals kinetic and quantitative differences that affect cell function. Sci. Signal. 11, eaat6753 (2018).

9. A. I. Salter, A. Rajan, J. J. Kennedy, R. G. Ivey, S. A. Shelby, I. Leung, M. L. Templeton, V. Muhunthan, V. Voillet, D. Sommermeyer, J. R. Whiteaker, R. Gottardo, S. L. Veatch, A. G. Paulovich, S. R. Riddell, Comparative analysis of TCR and CAR signaling informs CAR designs with superior antigen sensitivity and in vivo function. Sci. Signal. 14, eabe2606 (2021).

10. A. A. Griffith, K. P. Callahan, N. G. King, Q. Xiao, X. Su, A. R. Salomon, SILAC phosphoproteomics reveals unique signaling circuits in CAR-T cells and the inhibition of B cell-activating phosphorylation in target cells. J. Proteome Res. 21, 395–409 (2022).

11. V. M. Qin, C. D’Souza, P. J. Neeson, J. J. Zhu, Chimeric Antigen Receptor beyond CAR-T Cells. Cancers 13, 404 (2021).

12. R. G. Majzner, S. P. Rietberg, E. Sotillo, R. Dong, V. T. Vachharajani, L. Labanieh, J. H. Myklebust, M. Kadapakkam, E. W. Weber, A. M. Tousley, R. M. Richards, S. Heitzeneder, S. M. Nguyen, V. Wiebking, J. Theruvath, R. C. Lynn, P. Xu, A. R. Dunn, R. D. Vale, C. L. Mackall, Tuning the Antigen Density Requirement for CAR T-cell Activity. Cancer Discov. 10, 702–723 (2020).

13. S. Ramakrishna, S. L. Highfill, Z. Walsh, S. M. Nguyen, H. Lei, J. F. Shern, H. Qin, I. L. Kraft, M. Stetler-Stevenson, C. M. Yuan, J. D. Hwang, Y. Feng, Z. Zhu, D. Dimitrov, N. N. Shah, T. J. Fry, Modulation of Target Antigen Density Improves CAR T-cell Functionality and Persistence. Clin. Cancer Res. 25, 5329–5341 (2019).

14. X. He, Z. Feng, J. Ma, S. Ling, Y. Cao, B. Gurung, Y. Wu, B. W. Katona, K. P. O’Dwyer, D. L. Siegel, C. H. June, X. Hua, Bispecific and split CAR T cells targeting CD13 and TIM3 eradicate acute myeloid leukemia. Blood 135, 713–723 (2020).

15. S. Li, L. Shi, L. Zhao, Q. Guo, J. Li, Z. Liu, Z. Guo, Y. J. Cao, Split-design approach enhances the therapeutic efficacy of ligand-based CAR-T cells against multiple B-cell malignancies. Nat. Commun. 15, 9751 (2024).

16. S. Li, L. Shi, L. Zhao, Q. Guo, J. Li, Z. Liu, Z. Guo, Y. J. Cao, Split-design approach enhances the therapeutic efficacy of ligand-based CAR-T cells against multiple B-cell malignancies. Nat. Commun. 15, 9751 (2024).

17. B. I. Philipson, R. S. O’Connor, M. J. May, C. H. June, S. M. Albelda, M. C. Milone, 4-1BB costimulation promotes CAR T cell survival through noncanonical NF-κB signaling. Sci. Signal. 13, eaay8248 (2020).

18. B. Yao, X. Ye, Q. Kong, W. Chen, W. Li, C. Feng, A. He, G. Li, L. Chen, X. Chen, L. Hu, L. Xie, X. Qiu, X. Wang, Y. Lin, Y. Cao, J. Zhou, X. Zhang, H. Wang, R. Tian, Phosphotyrosine Signal Profiling of Clinical CAR-T Reveals Tonic Signaling Associated with Therapeutic Efficacy. bioRxiv, doi: 10.1101/2024.12.22.629940 (2024).

19. E. Hallumi, R. Shalah, W.-L. Lo, J. Corso, I. Oz, D. Beach, S. Wittman, A. Isenberg, M. Sela, H. Urlaub, A. Weiss, D. Yablonski, Itk Promotes the Integration of TCR and CD28 Costimulation through Its Direct Substrates SLP-76 and Gads. J. Immunol. 206, 2322–2337 (2021).

20. M. Barden, A. Holzinger, L. Velas, M. Mezősi-Csaplár, Á. Szöőr, G. Vereb, G. J. Schütz, A. A. Hombach, H. Abken, CAR and TCR form individual signaling synapses and do not cross-activate, however, can co-operate in T cell activation. Front. Immunol. 14 (2023).

21. K. M. Cappell, J. N. Kochenderfer, A comparison of chimeric antigen receptors containing CD28 versus 4-1BB costimulatory domains. Nat. Rev. Clin. Oncol. 18, 715–727 (2021).

22. Z. Zhao, M. Condomines, S. J. C. van der Stegen, F. Perna, C. C. Kloss, G. Gunset, J. Plotkin, M. Sadelain, Structural Design of Engineered Costimulation Determines Tumor Rejection Kinetics and Persistence of CAR T Cells. Cancer Cell 28, 415–428 (2015).

23. J. Rydzek, T. Nerreter, H. Peng, S. Jutz, J. Leitner, P. Steinberger, H. Einsele, C. Rader, M. Hudecek, Chimeric Antigen Receptor Library Screening Using a Novel NF-κB/NFAT Reporter Cell Platform. Mol. Ther. 27, 287–299 (2019).

24. K. S. Gordon, C. R. Perez, A. Garmilla, M. S. Y. Lam, J. J. Y. Aw, A. Datta, D. A. Lauffenburger, A. Pavesi, M. E. Birnbaum, Pooled screening for CAR function identifies novel IL-13Rα2-targeted CARs for treatment of glioblastoma. J. Immunother. Cancer 13, e009574 (2025).

25. D. Bloemberg, T. Nguyen, S. MacLean, A. Zafer, C. Gadoury, K. Gurnani, A. Chattopadhyay, J. Ash, J. Lippens, D. Harcus, M. Pagé, A. Fortin, R. A. Pon, R. Gilbert, A. Marcil, R. D. Weeratna, S. McComb, A High-Throughput Method for Characterizing Novel Chimeric Antigen Receptors in Jurkat Cells. Mol. Ther. Methods Clin. Dev. 16, 238–254 (2020).

26. K. S. Gordon, T. Kyung, C. R. Perez, P. V. Holec, A. Ramos, A. Q. Zhang, Y. Agarwal, Y. Liu, C. E. Koch, A. Starchenko, B. A. Joughin, D. A. Lauffenburger, D. J. Irvine, M. T. Hemann, M. E. Birnbaum, Screening for CD19-specific chimaeric antigen receptors with enhanced signalling via a barcoded library of intracellular domains. *Nat*. Biomed. Eng. 6, 855–866 (2022).

27. R. T. Abraham, A. Weiss, Jurkat T cells and development of the T-cell receptor signalling paradigm. Nat. Rev. Immunol. 4, 301–308 (2004).

28. F. Jahan, J. Koski, D. Schenkwein, S. Ylä-Herttuala, H. Göös, S. Huuskonen, M. Varjosalo, P. Maliniemi, J. Leitner, P. Steinberger, H.-J. Bühring, K. Vettenranta, M. Korhonen, Using the Jurkat reporter T cell line for evaluating the functionality of novel chimeric antigen receptors. *Front*. Mol. Med. 3 (2023).

29. R. Wan, B. Fu, X. Fu, Z. Liu, N. Simayi, Y. Fu, H. Liang, C. Li, W. Huang, SMAD4 Regulates the Expression of LCK Affecting Chimeric Antigen Receptor-T Cells Proliferation Through PI3K/Akt Signaling Pathway. J. Cell. Physiol. 240, e31520 (2025).

30. S. Sarntivijai, Y. Lin, Z. Xiang, T. F. Meehan, A. D. Diehl, U. D. Vempati, S. C. Schürer, C. Pang, J. Malone, H. Parkinson, Y. Liu, T. Takatsuki, K. Saijo, H. Masuya, Y. Nakamura, M. H. Brush, M. A. Haendel, J. Zheng, C. J. Stoeckert, B. Peters, C. J. Mungall, T. E. Carey, D. J. States, B. D. Athey, Y. He, CLO: The cell line ontology. J. Biomed. Semant. 5, 37 (2014).

31. F. L. Locke, A. Ghobadi, C. A. Jacobson, D. B. Miklos, L. J. Lekakis, O. O. Oluwole, Y. Lin, I. Braunschweig, B. T. Hill, J. M. Timmerman, others, Long-term safety and activity of axicabtagene ciloleucel in refractory large B-cell lymphoma (ZUMA-1): a single-arm, multicentre, phase 1–2 trial. Lancet Oncol. 20, 31–42 (2019).

32. A. Callahan, X. Zhang, A. Wang, A. Mojumdar, L. Zeng, X. Su, A. R. Salomon, CSF1R-CAR T cells induce CSF1R signalling and promote cancer cell growth. bioRxiv, 2024.12.17.629028 (2024).

33. L. Pillai-Kastoori, A. R. Schutz-Geschwender, J. A. Harford, A systematic approach to quantitative Western blot analysis. Anal. Biochem. 593, 113608 (2020).

34. A. Callahan, X. Y. Chua, A. A. Griffith, T. Hildebrandt, G. Fu, M. Hu, R. Wen, A. R. Salomon, Deep phosphotyrosine characterisation of primary murine T cells using broad spectrum optimisation of selective triggering. PROTEOMICS, 2400106 (2024).

35. K. Yu, A. R. Salomon, PeptideDepot: Flexible relational database for visual analysis of quantitative proteomic data and integration of existing protein information. PROTEOMICS 9, 5350–5358 (2009).

36. K. Yu, A. R. Salomon, HTAPP: High-throughput autonomous proteomic pipeline. PROTEOMICS 10, 2113–2122 (2010).

37. J. Belmont, T. Gu, A. Mudd, A. R. Salomon, A PLC-γ1 Feedback Pathway Regulates Lck Substrate Phosphorylation at the T-Cell Receptor and SLP-76 Complex. J. Proteome Res. 16, 2729–2742 (2017).

38. J. D. Storey, R. Tibshirani, Statistical significance for genomewide studies. Proc. Natl. Acad. Sci. 100, 9440–9445 (2003).

39. K. Krug, P. Mertins, B. Zhang, P. Hornbeck, R. Raju, R. Ahmad, M. Szucs, F. Mundt, D. Forestier, J. Jane-Valbuena, others, A curated resource for phosphosite-specific signature analysis*[S]. Mol. Cell. Proteomics 18, 576–593 (2019).

40. S. Rm. Clarke, M. Barnden, C. Kurts, F. R. Carbone, J. F. Miller, W. R. Heath, Characterization of the ovalbumin-specific TCR transgenic line OT-I: MHC elements for positive and negative selection. Immunol. Cell Biol. 78, 110–117 (2000).

41. X. Y. Chua, A. Salomon, Ovalbumin Antigen-Specific Activation of Human T Cell Receptor Closely Resembles Soluble Antibody Stimulation as Revealed by BOOST Phosphotyrosine Proteomics. J. Proteome Res. 20, 3330–3344 (2021).

42. W.-L. Lo, N. H. Shah, S. A. Rubin, W. Zhang, V. Horkova, I. R. Fallahee, O. Stepanek, L. I. Zon, J. Kuriyan, A. Weiss, Slow phosphorylation of a tyrosine residue in LAT optimizes T cell ligand discrimination. Nat. Immunol. 20, 1481–1493 (2019).

43. G. Gaud, R. Lesourne, P. E. Love, Regulatory mechanisms in T cell receptor signalling. Nat. Rev. Immunol. 18, 485–497 (2018).

44. S. Srivastava, S. R. Riddell, Engineering CAR-T cells: Design concepts. Trends Immunol. 36, 494–502 (2015).

45. O. Matalon, S. Fried, A. Ben-Shmuel, M. H. Pauker, N. Joseph, D. Keizer, M. Piterburg, M. Barda-Saad, Dephosphorylation of the adaptor LAT and phospholipase C–γ by SHP-1 inhibits natural killer cell cytotoxicity. Sci. Signal. 9, ra54–ra54 (2016).

46. B. Duan, D. Ye, S. Zhu, W. Jia, C. Lu, G. Wang, X. Guo, Y. Yu, C. Wu, J. Kang, HDAC10 promotes angiogenesis in endothelial cells through the PTPN22/ERK axis. Oncotarget Vol 8 No 37 (2017).

47. X. Zhang, Y. Yu, B. Bai, T. Wang, J. Zhao, N. Zhang, Y. Zhao, X. Wang, B. Wang, PTPN22 interacts with EB1 to regulate T-cell receptor signaling. FASEB J. 34, 8959–8974 (2020).

48. A. Ajina, J. Maher, Strategies to Address Chimeric Antigen Receptor Tonic Signaling. Mol. Cancer Ther. 17, 1795–1815 (2018).

49. J. H. Hanke, J. P. Gardner, R. L. Dow, P. S. Changelian, W. H. Brissette, E. J. Weringer, B. A. Pollok, P. A. Connelly, Discovery of a Novel, Potent, and Src Family-selective Tyrosine Kinase Inhibitor: STUDY OF Lck- AND FynT-DEPENDENT T CELL ACTIVATION (∗). J. Biol. Chem. 271, 695–701 (1996).

50. Y. A. Helou, V. Nguyen, S. P. Beik, A. R. Salomon, ERK Positive Feedback Regulates a Widespread Network of Tyrosine Phosphorylation Sites across Canonical T Cell Signaling and Actin Cytoskeletal Proteins in Jurkat T Cells. PLOS ONE 8, e69641 (2013).

51. L.-Y. Hsu, J. T. Rosenbaum, E. Verner, W. B. Jones, C. M. Hill, J. W. Janc, J. J. Buggy, R. D. Pawar, P. Ghosh, D. Li, N. Ding, J. C. Reneau, M. S. Khodadoust, Y. H. Kim, R. A. Wilcox, R. A. Miller, Synthesis and characterization of soquelitinib a selective ITK inhibitor that modulates tumor immunity. Npj Drug Discov. 1, 2 (2024).

52. A. M. Tousley, M. C. Rotiroti, L. Labanieh, L. W. Rysavy, W.-J. Kim, C. Lareau, E. Sotillo, E. W. Weber, S. P. Rietberg, G. N. Dalton, Y. Yin, D. Klysz, P. Xu, E. L. de la Serna, A. R. Dunn, A. T. Satpathy, C. L. Mackall, R. G. Majzner, Co-opting signalling molecules enables logic-gated control of CAR T cells. Nature 615, 507–516 (2023).

53. X. Shan, Czar, Michael J., Bunnell, Stephen C., Liu, Pinghu, Liu, Yusen, Schwartzberg, Pamela L., R. L. and Wange, Deficiency of PTEN in Jurkat T Cells Causes Constitutive Localization of Itk to the Plasma Membrane and Hyperresponsiveness to CD3 Stimulation. Mol. Cell. Biol. 20, 6945–6957 (2000).

54. X. Du, Darcy, Phillip K., Wiede, Florian, T. and Tiganis, Targeting Protein Tyrosine Phosphatase 22 Does Not Enhance the Efficacy of Chimeric Antigen Receptor T Cells in Solid Tumors. Mol. Cell. Biol. 42, e00449–21 (2022).

55. C. Petersen, M. Bell, H. Houke, Z. Yi, S. Gottschalk, G. Krenciute, IMMU-13. CRISPR/CAS9-MEDIATED SILENCING OF SHP-1 SIGNIFICANTLY ENHANCES THE ANTI-GLIOMA ACTIVITY OF IL-13Rα2 CAR T CELLS. Neuro-Oncol. 21, ii95– ii96 (2019).

56. C. Sun, P. Shou, H. Du, K. Hirabayashi, Y. Chen, L. E. Herring, S. Ahn, Y. Xu, K. Suzuki, G. Li, O. Tsahouridis, L. Su, B. Savoldo, G. Dotti, THEMIS-SHP1 Recruitment by 4-1BB Tunes LCK-Mediated Priming of Chimeric Antigen Receptor-Redirected T Cells. Cancer Cell 37, 216–225.e6 (2020).

57. F. Wiede, K. Lu, X. Du, S. Liang, K. Hochheiser, G. T. Dodd, P. K. Goh, C. Kearney, D. Meyran, P. A. Beavis, M. A. Henderson, S. L. Park, J. Waithman, S. Zhang, Z. Zhang, J. Oliaro, T. Gebhardt, P. K. Darcy, T. Tiganis, PTPN2 phosphatase deletion in T cells promotes anti-tumour immunity and CAR T-cell efficacy in solid tumours. EMBO J. 39, e103637 (2020).

58. O. Anannya, W. Huang, A. August, The kinase ITK controls a Ca2+-mediated switch that balances TH17 and Treg cell differentiation. Sci. Signal. 17, eadh2381.

