## Supplemental Material for "Inclusion of the ζ-chain drives phosphotyrosine signalling in CD19-CAR T cells"

**(A)** ζ-CAR

CD19 scFv CD8 tm ζ-sfGFP cyto

MALPVTALLLPLALLLHAARPDIQMTQTTSSLSASLGDRV
TISCRASQDISKYLNWYQQKPDGTVKLLIYHTSRLHSGVP
SRFSGSGSGTDYSLTISNLEQEDIATYFCQQGNTLPYTFG
GGTKLEITGGGGSGGGGSGGGGSEVKLQESGPGLVAPSQS
LSVTCTVSGVSLPDYGVSWIRQPPRKGLEWLGVIWGSETT
YYNSALKSRLTIIKDNSKSQVFLKMNSLQTDDTAIYYCAK
HYYYGGSYAMDYWGQGTSVTVSSGSGSEQKLISEEDLGST
TTPAPRPPTPAPTIASQPLSLRPEACRPAAGGAVHTRGLD
FACDIYIWAPLAGTCGVLLLSLVITLYCSRRVKFSRSADA
PAYQQGQNQLYNELNLGRREEYDVLDKRRGRDPEMGGKPQ
RRKNPQEGLYNELQKDKMAEAYSEIGMKGERRRGKGHDGL
YQGLSTATKDTYDALHMQALPPRTSGSGSMSKGEELFTGV
VPILVELDGDVNGHKFSVRGEGEGDATNGKLTLKFICTTG
KLPVPWPTLVTTLTYGVQCFSRYPDHMKRHDFFKSAMPEG
YVQERTISFKDDGTYKTRAEVKFEGDTLVNRIELKGIDFK
EDGNILGHKLEYNFNSHNVYITADKQKNGIKANFKIRHNV
EDGSVQLADHYQQNTPIGDGPVLLPDNHYLSTQSVLSKDP
NEKRDHMVLLEFVTAAGITHGMDELYK

FMC63 scFv
CD8α tm
TCRζ
sfGFP

**(B)** 28ζ-CAR

CD19 scFv CD8 tm CD28-ζ-sfGFP cyto

MALPVTALLLPLALLLHAARPDIQMTQTTSSLSASLGDRV
TISCRASQDISKYLNWYQQKPDGTVKLLIYHTSRLHSGVP
SRFSGSGSGTDYSLTISNLEQEDIATYFCQQGNTLPYTFG
GGTKLEITGGGGSGGGGSGGGGSEVKLQESGPGLVAPSQS
LSVTCTVSGVSLPDYGVSWIRQPPRKGLEWLGVIWGSETT
YYNSALKSRLTIIKDNSKSQVFLKMNSLQTDDTAIYYCAK
HYYYGGSYAMDYWGQGTSVTVSSGSGSEQKLISEEDLGST
TTPAPRPPTPAPTIASQPLSLRPEACRPAAGGAVHTRGLD
FACDIYIWAPLAGTCGVLLLSLVITLYCSRSKRSRLLHSD
YMNMTPRRPGPTRKHYQPYAPPRDFAAYRSRVKFSRSADA
PAYQQGQNQLYNELNLGRREEYDVLDKRRGRDPEMGGKPQ
RRKNPQEGLYNELQKDKMAEAYSEIGMKGERRRGKGHDGL
YQGLSTATKDTYDALHMQALPPRTSGSGSMSKGEELFTGV
VPILVELDGDVNGHKFSVRGEGEGDATNGKLTLKFICTTG
KLPVPWPTLVTTLTYGVQCFSRYPDHMKRHDFFKSAMPEG
YVQERTISFKDDGTYKTRAEVKFEGDTLVNRIELKGIDFK
EDGNILGHKLEYNFNSHNVYITADKQKNGIKANFKIRHNV
EDGSVQLADHYQQNTPIGDGPVLLPDNHYLSTQSVLSKDP
NEKRDHMVLLEFVTAAGITHGMDELYK

FMC63 scFv
CD8α tm
CD28 cyto
TCRζ
sfGFP

**(C)** BBζ-CAR

CD19 scFv CD8 tm 41BB-ζ-sfGFP cyto

MALPVTALLLPLALLLHAARPDIQMTQTTSSLSASLGDRV
TISCRASQDISKYLNWYQQKPDGTVKLLIYHTSRLHSGVP
SRFSGSGSGTDYSLTISNLEQEDIATYFCQQGNTLPYTFG
GGTKLEITGGGGSGGGGSGGGGSEVKLQESGPGLVAPSQS
LSVTCTVSGVSLPDYGVSWIRQPPRKGLEWLGVIWGSETT
YYNSALKSRLTIIKDNSKSQVFLKMNSLQTDDTAIYYCAK
HYYYGGSYAMDYWGQGTSVTVSSGSGSEQKLISEEDLGST
TTPAPRPPTPAPTIASQPLSLRPEACRPAAGGAVHTRGLD
FACDIYIWAPLAGTCGVLLLSLVITLYCSRKRGRKKLLYI
FKQPFMRPVQTTQEEDGCSCRFPEEEEGGCELRVKFSRSA
DAPAYQQGQNQLYNELNLGRREEYDVLDKRRGRDPEMGGK
PQRRKNPQEGLYNELQKDKMAEAYSEIGMKGERRRGKGHD
GLYQGLSTATKDTYDALHMQALPPRTSGSGSMSKGEELFT
GVVPILVELDGDVNGHKFSVRGEGEGDATNGKLTLKFICT
TGKLPVPWPTLVTTLTYGVQCFSRYPDHMKRHDFFKSAMP
EGYVQERTISFKDDGTYKTRAEVKFEGDTLVNRIELKGID
FKEDGNILGHKLEYNFNSHNVYITADKQKNGIKANFKIRH
NVEDGSVQLADHYQQNTPIGDGPVLLPDNHYLSTQSVLSK
DPNEKRDHMVLLEFVTAAGITHGMDELYK

FMC63 scFv
CD8α tm
41BB cyto
TCRζ
sfGFP

**(D)** 28BBζ-CAR

CD19 scFv CD8 tm CD28-41BB-ζ-sfGFP cyto

MALPVTALLLPLALLLHAARPDIQMTQTTSSLSASLGDRV
TISCRASQDISKYLNWYQQKPDGTVKLLIYHTSRLHSGVP
SRFSGSGSGTDYSLTISNLEQEDIATYFCQQGNTLPYTFG
GGTKLEITGGGGSGGGGSGGGGSEVKLQESGPGLVAPSQS
LSVTCTVSGVSLPDYGVSWIRQPPRKGLEWLGVIWGSETT
YYNSALKSRLTIIKDNSKSQVFLKMNSLQTDDTAIYYCAK
HYYYGGSYAMDYWGQGTSVTVSSGSGSEQKLISEEDLGST
TTPAPRPPTPAPTIASQPLSLRPEACRPAAGGAVHTRGLD
FACDIYIWAPLAGTCGVLLLSLVITLYCSRSKRSRLLHSD
YMNMTPRRPGPTRKHYQPYAPPRDFAAYRSKRGRKKLLYI
FKQPFMRPVQTTQEEDGCSCRFPEEEEGGCELRVKFSRSA
DAPAYQQGQNQLYNELNLGRREEYDVLDKRRGRDPEMGGK
PQRRKNPQEGLYNELQKDKMAEAYSEIGMKGERRRGKGHD
GLYQGLSTATKDTYDALHMQALPPRTSGSGSMSKGEELFT
GVVPILVELDGDVNGHKFSVRGEGEGDATNGKLTLKFICT
TGKLPVPWPTLVTTLTYGVQCFSRYPDHMKRHDFFKSAMP
EGYVQERTISFKDDGTYKTRAEVKFEGDTLVNRIELKGID
FKEDGNILGHKLEYNFNSHNVYITADKQKNGIKANFKIRH
NVEDGSVQLADHYQQNTPIGDGPVLLPDNHYLSTQSVLSK
DPNEKRDHMVLLEFVTAAGITHGMDELYK

FMC63 scFv
CD8α tm
CD28 cyto
41BB cyto
TCRζ
sfGFP

**Supporting Figure 1:** Amino acid sequences for (A) ζ-CAR, (B) 28ζ-CAR, (C) BBζ-CAR, and (D) 28BBζ-CAR.

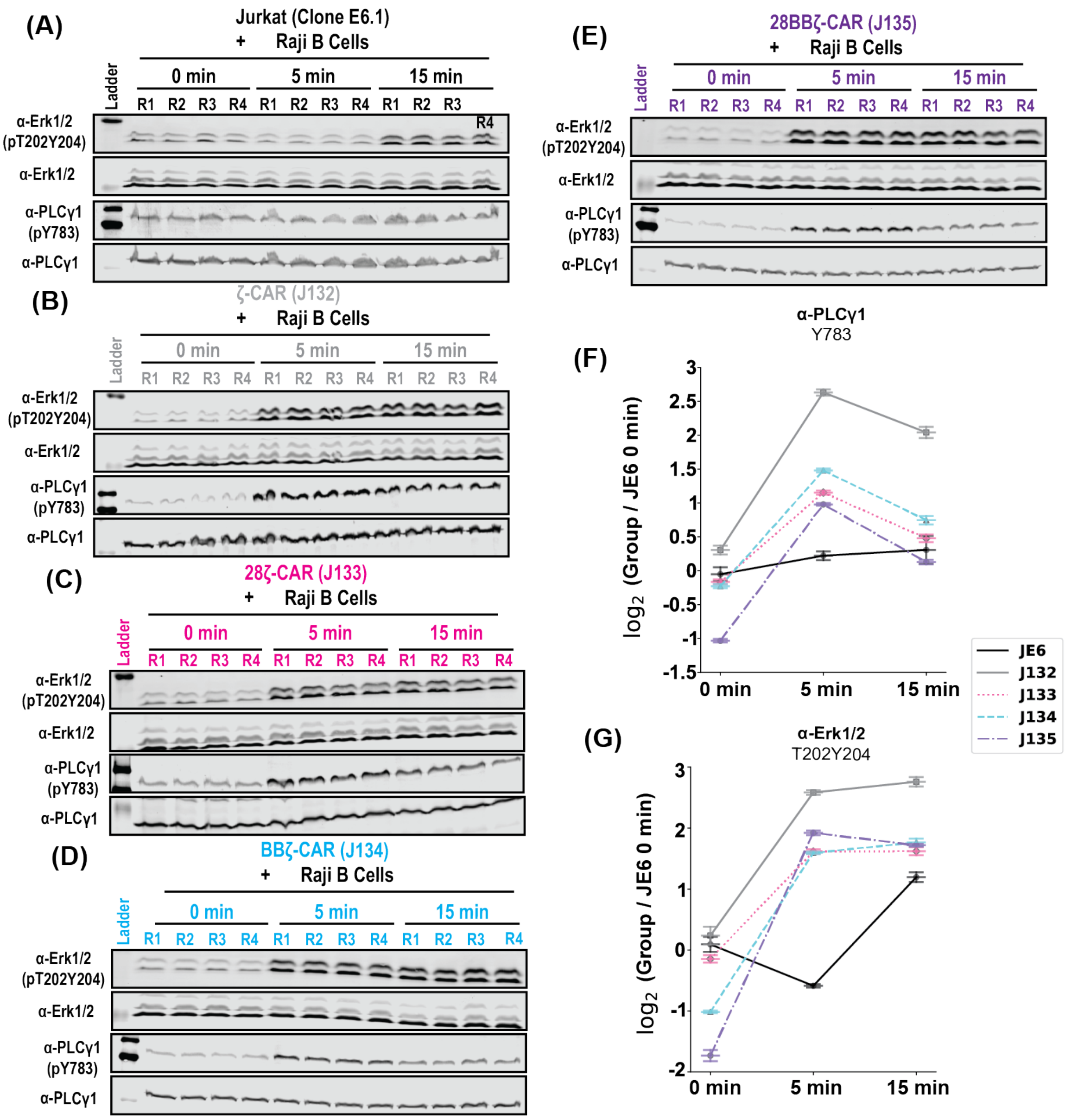


**Supporting Figure 2:** Co-culture stimulation experiments show similar induction of pY sites on TCR signalling proteins. (A) Western blot analysis of Erk1$^{T202Y204}$/Erk2$^{T185Y187}$, Zap70$^{Y493}$, and PLCγ1$^{Y783}$ in co-cultures of Jurkat T cells and Raji B cells. (B-E) As in A, except for co-cultures of Raji B cells and ζ-CAR, 28ζ-CAR, BBζ-CAR, and 28BBζ-CAR, respectively. (F, G) Quantification of PLC1$^{Y783}$ and Erk1$^{T202Y204}$/Erk2$^{T185Y187}$ phosphorylation, respectively. All groups are relative to the JE6 0m group. Supporting Table 1 contains the results of Fisher's LSD with the Holm-Sidak correction.

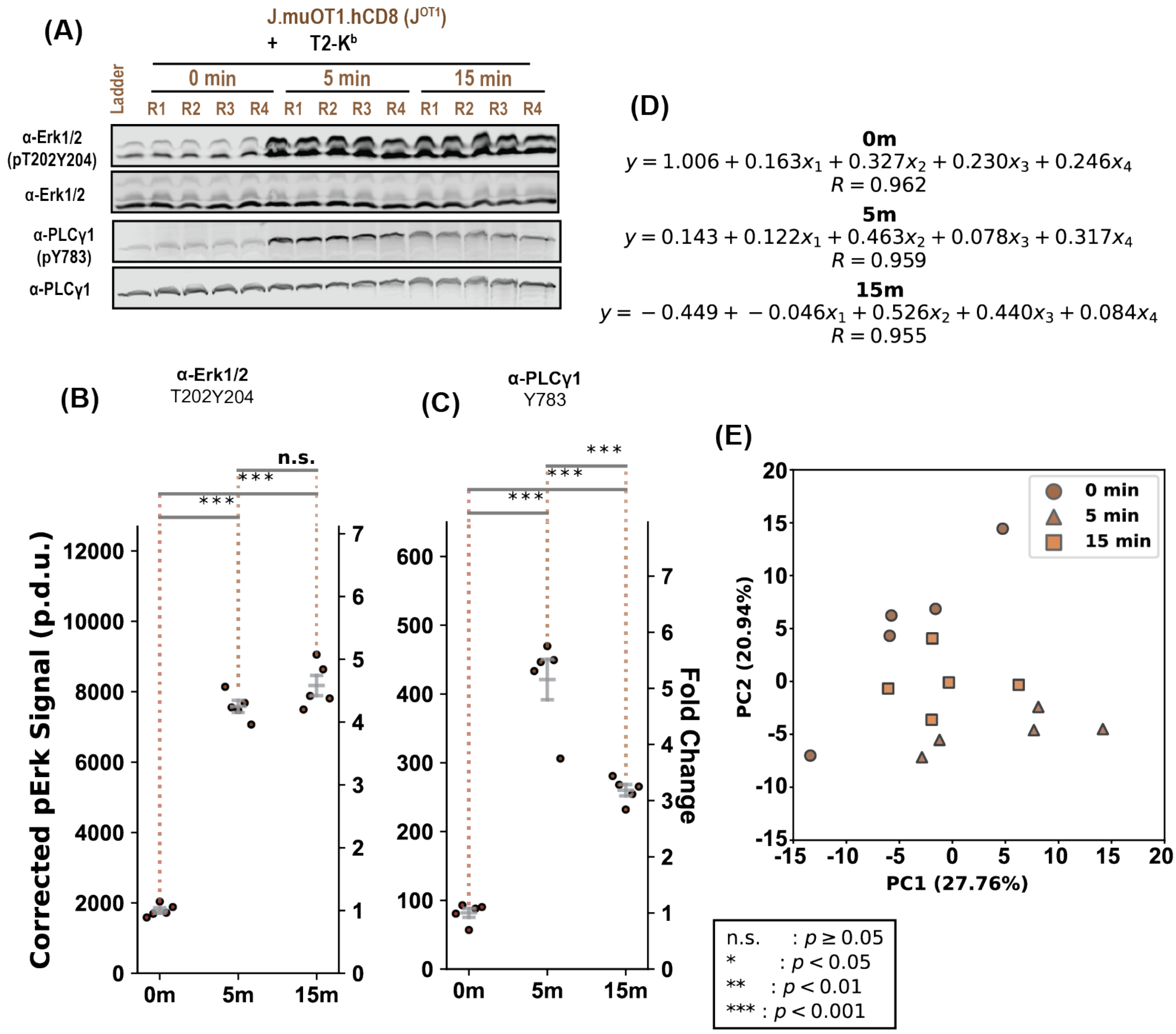


**Supporting Figure 3:** Co-culture of Jurkat cells expressing the murine OT1 transgenic TCR and human CD8 with OVA$^{257\text{-}264}$ loaded T2-K$^{b}$ cells induces highly reproducible TCR stimulation. (A) Western blot analysis of Erk1$^{T202Y204}$/Erk2$^{T185Y187}$ and PLCγ1$^{Y783}$ in co-cultures of J$^{OT1}$ cells and pT2-K$^{b}$ cells. (B, C) Quantification of phosphorylation of Erk1$^{T202Y204}$/Erk2$^{T185Y187}$ and PLCγ1$^{Y783}$, respectively, in J$^{OT1}$ cell and pT2-K$^{b}$ cell co-cultures. Supporting Table 1 contains the results of Fisher's LSD with the Holm-Sidak correction. (D) Multiple linear regression on pY sites measured in all replicates during co-culture of J$^{OT1}$ and pT2-K$^{b}$. (E) Principal component analysis of pY sites measured in all replicates of J$^{OT1}$ cells during co-culture with pT2-K$^{b}$ cells.

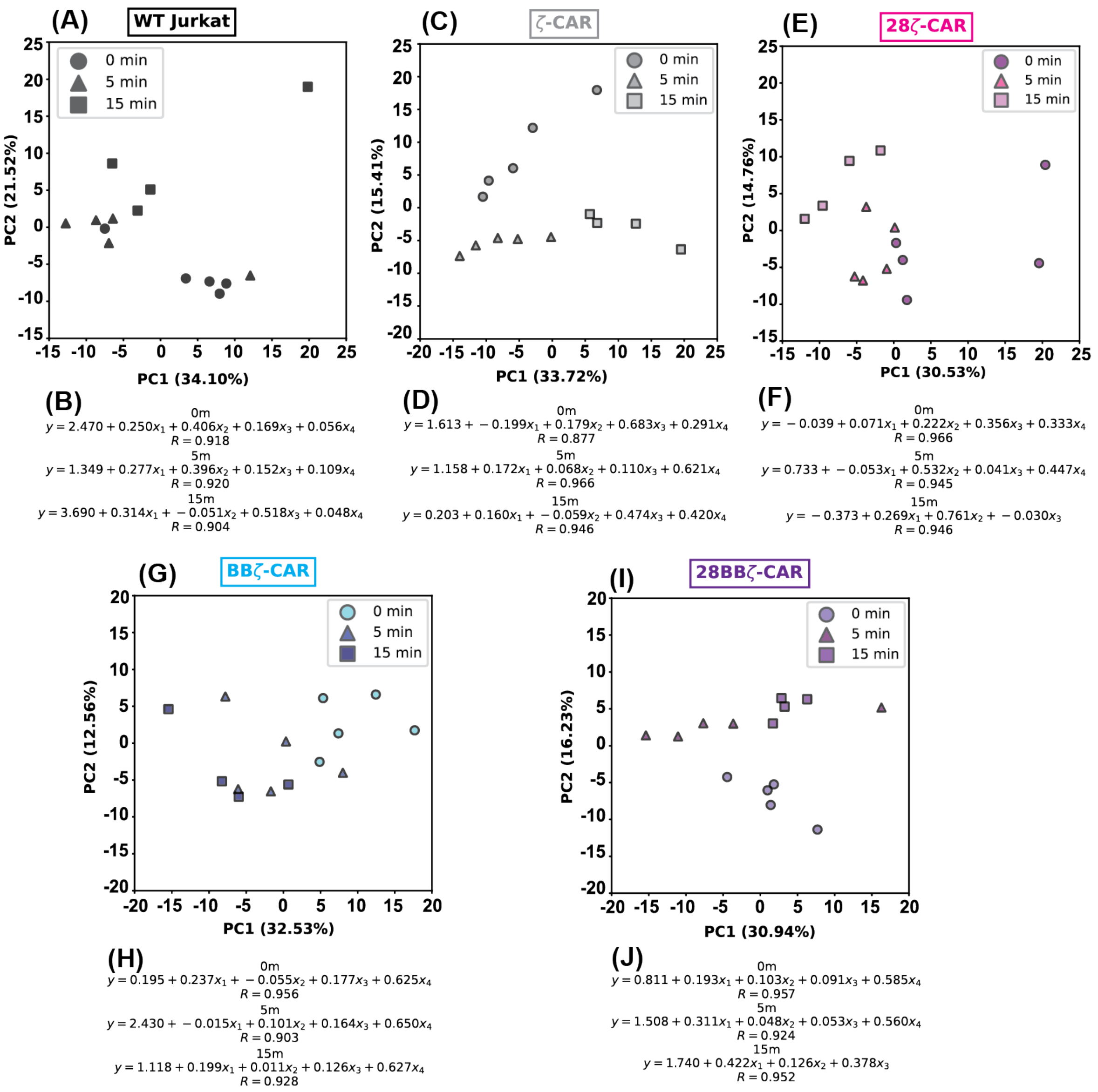


**Supporting Figure 4:** pY proteomics data for CD19-CAR T cells is highly reproducible. (A) Principal component analysis of pY sites measured in all replicates of WT Jurkat cells during co-culture with Raji B cells. (B) Multiple linear regression on pY sites measured in all replicates during co-culture of WT Jurkat cells with Raji B cells. (C-D) As in A & B, except for ζ-CAR. (E-F) As in A & B, except for 28ζ-CAR. (G-H) As in A & B, except for BBζ-CAR. (I-J) As in A & B, except for 28BBζ-CAR.

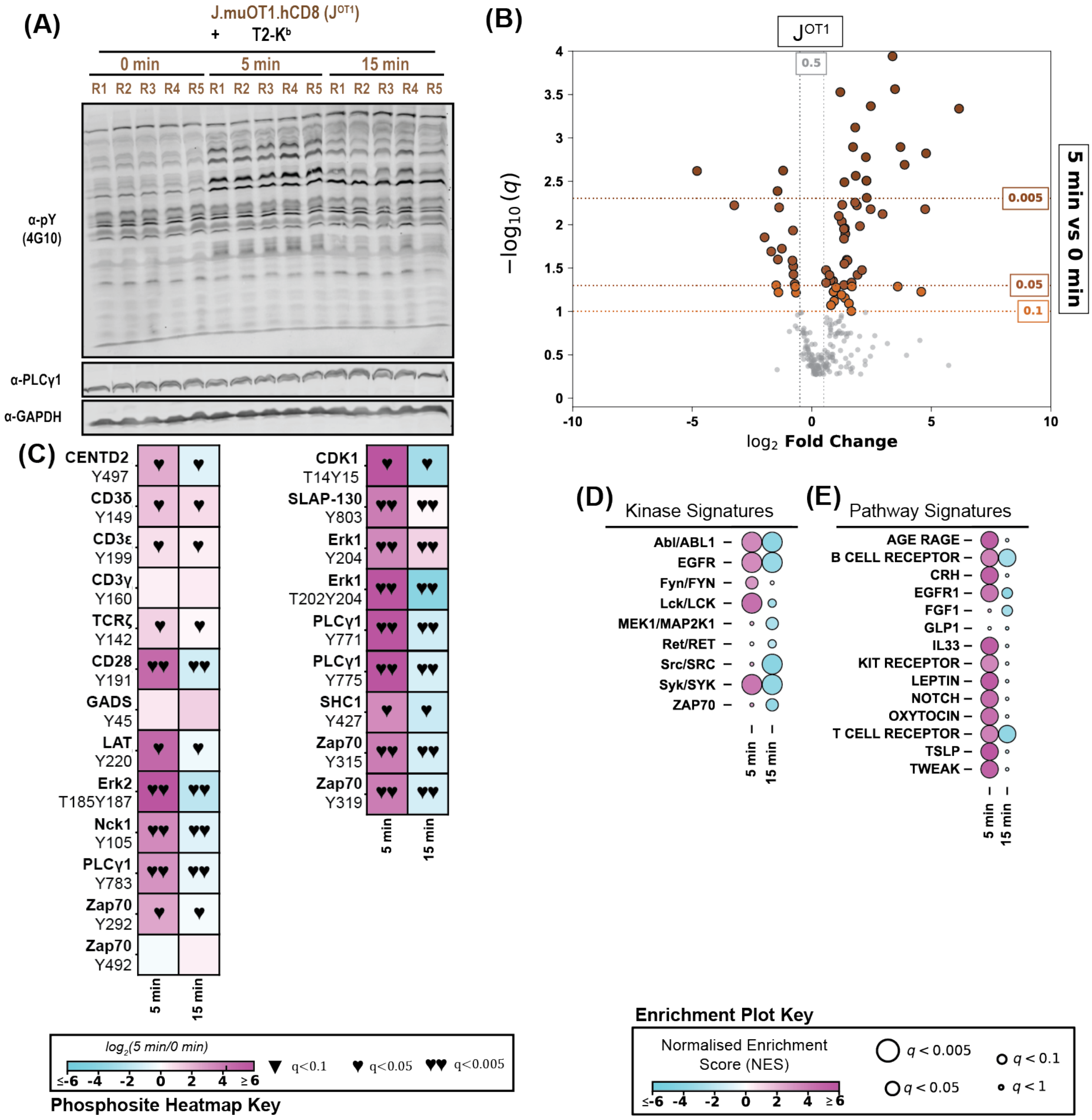


**Supporting Figure 5:** pT2-K$^b$ cells induce strong TCR pY signalling in J$^{OT1}$ cells. (A) Western blot analysis of global tyrosine phosphorylation in pY proteomics samples during a time course co-culture of J$^{OT1}$ cells and pT2-K$^b$ cells. (B) Volcano plot analysis showing significance and fold-change of pY sites after 5 minutes of J$^{OT1}$ and pT2-K$^b$ co-culture. (C) Heat maps comparing 5 minutes or 15 minutes with 0 minutes of co-culture between pT2-K$^b$ cells and J$^{OT1}$ cells. Sites presented are a selection of known TCR-responsive pY sites. ▼ indicates q < 0.1 (trending towards significance), ♥ indicates q < 0.05, and ♥♥ indicates q < 0.005. (D-E) Post translational modification signature enrichment analysis using pY site data showing enrichment of PhosphoSitePlus kinase substrate signatures and NetPath pathway signatures, respectively, for co-culture between pT2-K$^b$ cells and J$^{OT1}$ cells.

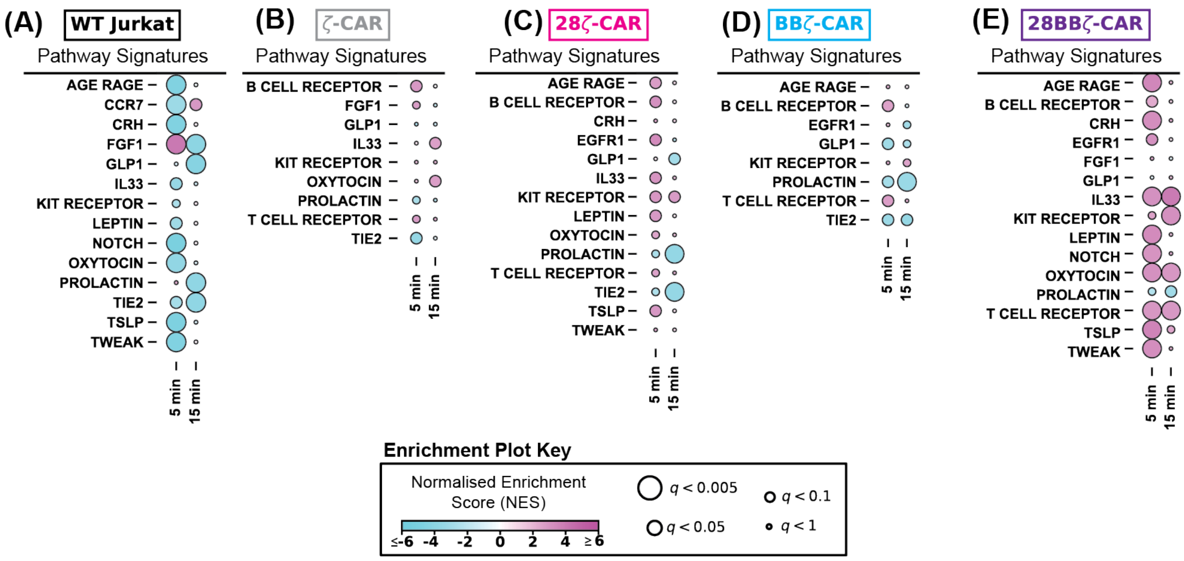


**Supporting Figure 6:** PTM-SEA reveals high engagement of signalling cascades in the third generation 28BBζ-CAR. (A-E) Post translational modification signature enrichment analysis using pY site data showing enrichment of NetPath pathway signatures for WT Jurkats, ζ-CAR, 28ζ-CAR, BBζ-CAR, and 28BBζ-CAR, respectively.

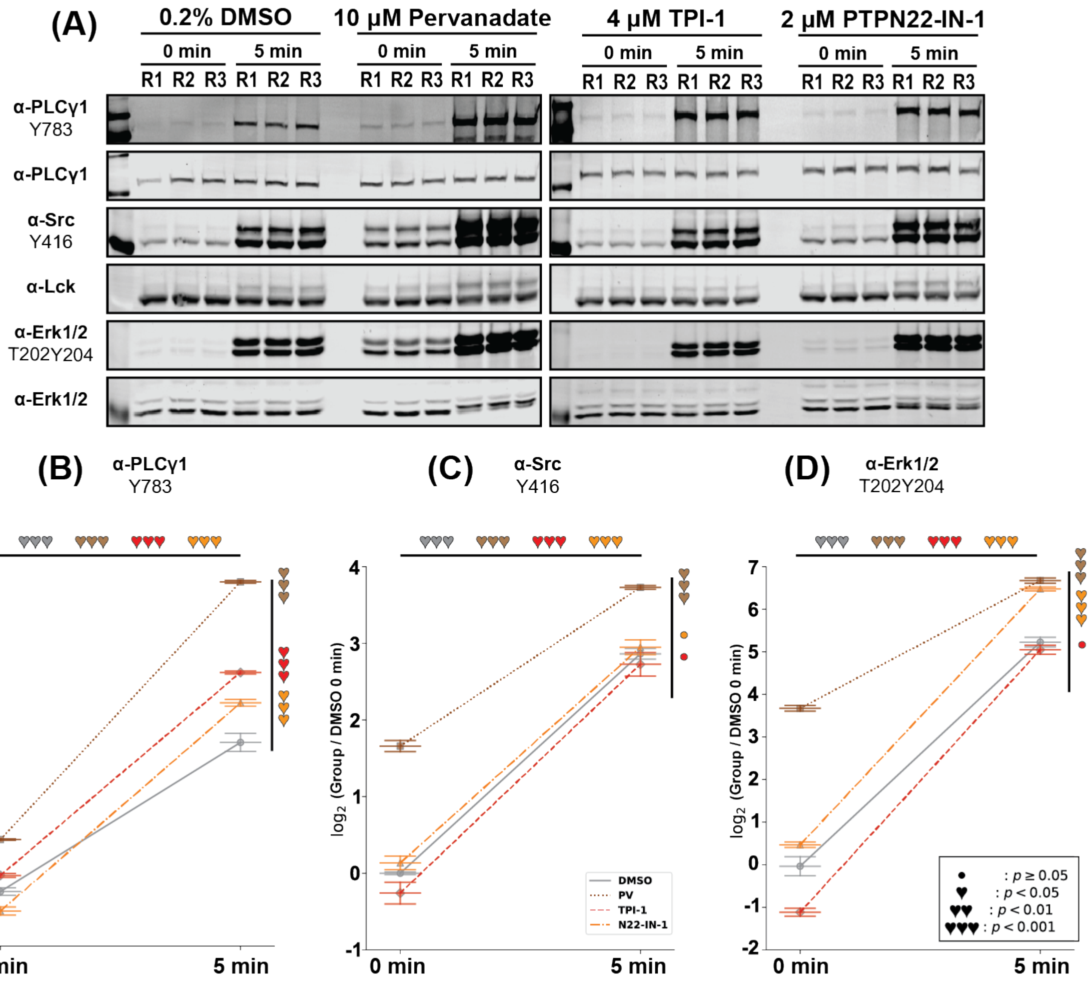

**Supporting Figure 7:** Effects of phosphatase inhibitors Pervanadate (global), TPI-1 (SHP-1), and PTPN22-IN-1 (PTPN22) on pY induction of T cell signalling proteins after CD3ε/CD28 antibody crosslinking. (A) Western blot analysis of PLC1$^{Y783}$, Lck$^{Y394}$, and Erk1$^{T202Y204}$/Erk2$^{T185Y187}$ phosphorylation in the basal and stimulated states of untreated or phosphatase inhibitor treated Jurkat T cells. (B-D) Quantification of PLC1$^{Y783}$, Lck$^{Y394}$, and Erk1$^{T202Y204}$/Erk2$^{T185Y187}$ phosphorylation, respectively. Supporting Table 9 contains the results of Fisher's LSD with the Holm-Sidak correction.

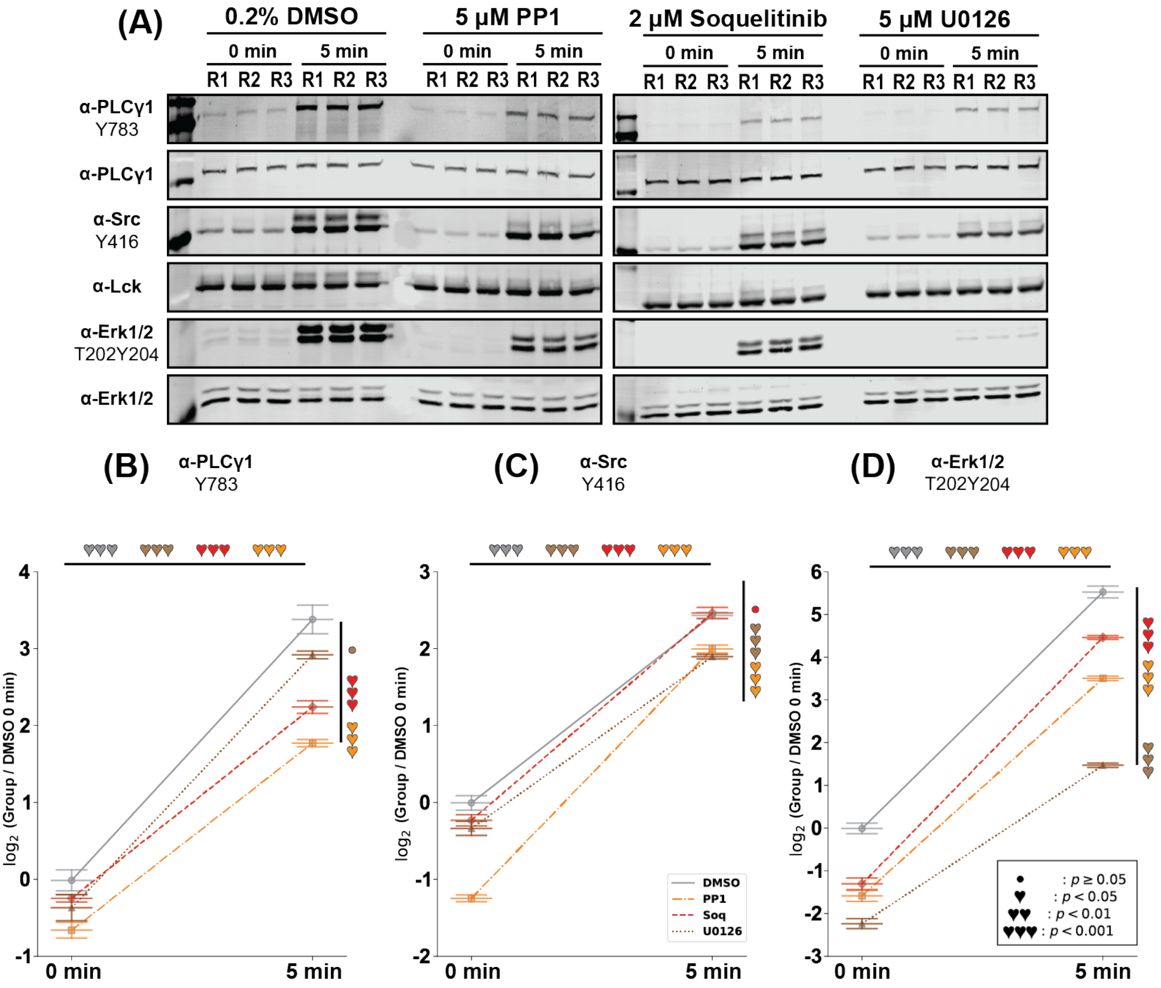


**Supporting Figure 8:** Effects of kinase inhibitors PP1 (Lck), Soquelitinib (Itk), and U0126 (MEK1/2) on pY induction of T cell signalling proteins after CD3ε/CD28 antibody crosslinking. (A) Western blot analysis of PLC1$^{Y783}$, Lck$^{Y394}$, and Erk1$^{T202Y204}$/Erk2$^{T185Y187}$ phosphorylation in the basal and stimulated states of untreated or kinase inhibitor treated Jurkat T cells. (B-D) Quantification of PLC1$^{Y783}$, Lck$^{Y394}$, and Erk1$^{T202Y204}$/Erk2$^{T185Y187}$ phosphorylation, respectively. Supporting Table 10 contains the results of Fisher's LSD with the Holm-Sidak correction.
